# Urinary proteome of dogs with kidney injury during babesiosis

**DOI:** 10.1101/516120

**Authors:** D. Winiarczyk, K. Michalak, L. Adaszek, M. Winiarczyk, J. Madany, S. Winiarczyk

## Abstract

This study aimed to identify proteins found in the urine of dogs with renal dysfunction leading to acute injury during the natural course of babesiosis (n=10) and to compare them with proteins of a control group (n=10) to reveal any potential biomarkers of renal damage. Pooled urine samples of both groups were separated by 2D electrophoresis (two dimensional electrophoresis), followed by the identification of all proteins using MALDI-TOF mass spectrometry (matrix assisted laser desorption ionization-time of flight). In total, 176 proteins were identified in the urine samples from healthy dogs, and 403 proteins were identified in the urine samples from dogs with babesiosis. Of the 176 proteins, 146 were assigned exclusively to healthy dogs, and 373 of the 403 proteins were assigned exclusively to dogs with babesiosis; 30 proteins were common to both groups. Characteristic analysis of the 373 proteins found in dogs with babesiosis led to the isolation of 8 proteins associated with 10 metabolic pathways that were attributed to immune and inflammatory response development. Furthermore, it was hypothesized that the epithelial-mesenchymal transition might play an important role in mechanisms underlying pathological renal tissue changes during babesiosis, as indicated by a causal relationship network built by combining 5 of the 10 selected metabolic pathways and 4 of the 8 proteins associated with these pathways. These included cadherins, gonadotropin releasing hormone receptors, inflammatory responses mediated by chemokine and cytokine signalling pathways, integrins, interleukin and TGF-β (transforming growth factor β) pathways. These pathways were linked by interleukin-13, bone morphogenetic protein 7, α2(1) collagen, and FER tyrosine kinase, which are potential damage biomarkers during babesiosis in dogs that might be assigned to early renal injury.

## Introduction

After heart failure, kidney disease is the most frequent cause of lowering the quality and shortening the life of people and dogs. Kidney injuries, which take one of two forms, acute kidney injury (AKI) and chronic kidney disease (CKD), are caused by various factors. In humans, 7.8% of patients with AKI also develop CKD, and 4.9% of patients reach end-stage renal disease [1]. Most AKI cases in medicine and veterinary science are diagnosed based on the serum or plasma concentrations of non-protein nitrogenous creatinine (Cr) and urea compounds. The sensitivity of this method is small and not suitable for early AKI detection [2], and seeking markers and methods adequate for the early detection of glomeruli and/or tubule injury before the decreased glomerular filtration rate (GFR) is signalled by increased Cr concentrations is thus necessary [3–6]. One such method is proteomic analysis, which compares the protein profiles of normal urine with those typical for a given disease to select potential diagnostic, therapeutic and prognostic biomarkers [7,8]. With decreased GMRs and subsequent azotemia and urea, AKI is among the most frequently occurring complications of babesiosis in dogs and may provide a natural model for identifying early and specific markers of kidney injury in this species [9,10]. Moreover, during babesiosis occurring naturally in dogs, AKI potentially provides a good model for selected studies on AKI in humans. This is indicated by comparative analysis of the urine proteomes in humans and dogs, as many proteins related to human diseases, including kidney diseases, have been identified in canine urine [11,12]. In addition, domestic dogs *(Canis lupus familiaris)* are increasingly perceived as an excellent animal model for studying complex human diseases [13]. Because they have a fully described genome and share a habitat with humans, dogs may be used for epidemiological studies on diseases shared between the two species. Canine DNA and protein sequences are much closer to humans than those of mice, suggesting that dogs are also more similar to many aspects of human biology than mice [14–16]. This study aimed to identify proteins found in the urine of dogs with renal dysfunction leading to acute injury during the natural course of babesiosis and compare them with proteins of the control group to reveal any potential biomarkers of renal injury.

## Materials and methods

### Animals and sample collection

Dogs were enrolled during routine admission to Faculty of Veterinary Medicine clinics at the University of Life Sciences in Lublin. Informed consent was obtained from the owners prior to the clinical investigations and sample collections. The studies were reviewed and approved by the Ethics Committee of the University of Life Sciences in Lublin (Poland) No 70/2018. The study involved 20 mixed-breed dogs (10 males, 10 females) weighing 5–8 kg (median 6.2 kg) and aged 2–7 years (median 4.35 years), divided into two groups. All dogs underwent individual clinical and laboratory tests to determine their health status, and in particular in diseased group to show signs of kidney damage. Group 1 (study group, n=10; five males and five females), consisted of dogs naturally infected with *B. canis*, while group 2 (control group, n=10; five males and five females) consisted of healthy [17]. All dogs in the first group showed symptoms of babesiosis (apathy, anorexia, changes in urine colour, pale mucous membranes), and haematology analysis revealed thrombocytopenia (platelets 12–88 × 10^9^/l) and anaemia (erythrocytes 3.5–5.3 × 10^12^/l.) All dogs were nonazotemic, serum creatinine concentration remained within the reference range. All dogs in this group had *Babesia*-positive blood smears, which were additionally confirmed by PCR, performed according to the protocol described by Adaszek and Winiarczyk [9]. Possible co-infections (borreliosis, anaplasmosis, ehrlichiosis) were excluded in all dogs based on PCR and ELISA results [18]. All dogs of the first group were successfully treated with imidocarb (5 mg/kg s.c.). Dogs of group 2 were clinically healthy and were referred to the clinic for vaccination purposes. Blood smear analysis and PCR for *B. canis* gave negative results for all animals of this group. Voided midstream urine samples were collected in the morning, and each sample was centrifuged on the day of collection at 500 × g for 10 minutes at 4°C. The supernatants were removed, and protease inhibitors were added (Protease Inhibitor Cocktail, Roche Diagnostic Corp.). Urine protein (low proteinuria denoted by “+”, moderate proteinuria denoted by “++”, and severe proteinuria denoted by “+++”) and Cr concentrations were measured by the enzymatic colorimetric method (BS-130 analyser, Mindray), and basic urinalysis with microscopic sediment analysis was performed on the fresh urine samples. Urine specific gravity (USG) was measured using a refractometer. The remaining urine was frozen at –80°C for further analysis. Macroscopic evaluation of urine in group 1 showed yellow to dark brown sample colours, while all group 2 samples were yellow. Urine protein analysis revealed proteinuria in eight of the 10 group 1 dogs, and eight dogs of this group also had urine protein/Cr ratios > 0.5. Urine dipstick analysis showed haemoglobinuria in seven of the 10 group 1 dogs, which was severe (+++) in two dogs. Urine specific gravity decreased in all diseased dogs and amounted to 1.015 on average. None of the control group dogs had proteinuria or haemoglobinuria. Statistically higher concentrations of urinary biomarkers (uIgG/uCr, uTHP/uCr, and uRBP/uCr) were found in the urine samples of all dogs with babesiosis compared to those in the control animals (p < 0.05), indicating dysfunctional glomerular and tubular kidney regions [17]. For proteomic analysis, 10 individual urine samples (0.5 ml each) from groups 1 and 2 were collected and pooled from affected and healthy dogs, respectively. Each pooled urine sample was subjected to desaltation on the filter to enable quick ultrafiltration with a high-density coefficient (Amicon Ultra Merck). Protein concentrations were measured with a microlitre spectrophotometer (NANO), and the urine samples were then prepared and subjected to 2D electrophoresis. Each individual gel spot was then analysed by mass spectrometry with the MALDI-TOF (matrix-assisted laser desorption ionization – time of flight) technique.

### 2D electrophoresis

Two-dimensional electrophoresis was used to separate the proteins contained in the tested urine samples [19]. Preliminary tests have shown that the optimum amount of protein for 2D electrophoresis is 85 μg; thus, this amount of protein was broken down via a precipitation and purification kit (ReadyPrep™ 2-D Cleanup Kit, Bio-Rad, Warsaw, Poland). The obtained protein pellets were then dissolved in a rehydration buffer, and the resulting solutions were applied to a rehydration plate and covered with 17-cm immobilized pH gradient (IPG-immobilized pH gradient) strips for isoelectric focusing (pH 3-10, Bio-Rad). To soak the gel present on the strips with the protein sample, the strips were removed after a 12-hour rehydration period and then subjected to the first electrophoresis dimension (IEF-100 Hoefer; 250 V/30 min; 10 000 V/3 hrs; 60 kV/hr, with a current limit of 50 μA/strip hrs). Under the influence of the electric field, proteins contained in the strips were subjected to migration by siting in a location corresponding to the isoelectric point of the given protein. After separation, the IPG strips were prepared for the second electrophoresis dimension to separate the proteins by molecular mass. Vertical electrophoretic separation utilized 12.5% polyacrylamide gels and the following current parameters: 600 V/30 mA/100 W in an electrophoretic chamber (PROTEAN® II xi, Bio-Rad). The obtained gels were subjected to a standard colouring procedure with silver in the presence of formaldehyde as a regulator. The protein spots were cut out of the gels, decolourised, reduced and alkylated using dithiothreitol and iodoacetamide [20]. Gel fragments containing proteins were subjected to digestion to obtain shorter peptide fragments. Trypsin digestion occurred in 50 mM ammonium bicarbonate buffer at 37°C for 12 hours (Promega, Trypsin Gold, Mass Spectrometry Grade, Technical Bulletin) [21]. The obtained peptides were subsequently eluted from the gel with a water/acetonitrile/TFA solution (v:v 450:500:50). The extracted peptides were purified using C18 Zip-TIP pipette tips according to the manufacturer’s instructions (Merck Chemicals, Billerica, MA, USA, PR 02358, Technical Note) and applied to the MTP AnchorChip 384 plate (Bruker, Bremen, Germany).

### Mass spectrometry

After the protein samples were dried on the MPT AnchorChip 384 plate, their surfaces were covered with a super-saturated solution of α-cyano4-hydroxycinnamic acid (HCCA, Bruker), functioning as a matrix mediating the transmission of energy to the sample. Simultaneously, 0.5 μl of a peptide standard was applied to the calibration fields (Peptide Calibration Standard II, Bruker), which were also covered with the matrix solution. Spectrometric analysis was performed using the Ultraflextreme III MALDI TOF/TOF (Bruker), and flexControl 3.3 (Bruker) software was applied for mass spectra collection. The obtained peptides were subjected to mild ionization using the MALDI-TOF instrument in the linear mode within the 900-4000 Da mass scope in the reflectron mode. The obtained mass spectra were analysed with flexAnalysis 3.4 (Bruker) software as follows: smoothing (Savitsky-Golay method), baseline subtraction (Top Hat baseline algorithm), and peak geometry (Stanford Network Analysis Platform (SNAP) algorithm). All peaks with signal to noise ratios > 3 were qualified for further analysis. Experimental data were analysed using the abovementioned software to exclude peaks originating from trypsin or environmental pollution. To ensure correct identification, selecting possible post-translation modifications using BioTools 3.2 (Bruker) software was essential. Post-translation modifications were derived from both the methodology used as well as the metabolic processes of the patients. The obtained spectra were compared to the Swiss-Prot database restricted to “bony vertebrate” taxa using Mascot 2.2 software with a maximum error of 0.3 Da. If the obtained result was not statistically significant, the original peptide ions were subjected to fragmentation in the tandem spectrometry mode [22,23].

## Results

Based on the clinicopathological variables all dogs with babesiosis met the criteria for early phase of AKI. They had proteinuria with UPC>0.5, decreased urine specific gravity amounted to 1.015 on average and significantly elevated value of uIgG/uCr, uTHP/uCr, and uRBP/uCr that indicated glomerular and tubular damage.

In this study, 176 proteins were identified in pooled urine samples collected from healthy dogs, and 403 proteins were identified in pooled urine samples collected from dogs with babesiosis. Tables 1 and 2 contain lists of the proteins, along with their names, scores, molecular weights, number of matches, UniProt base accession numbers and hyperlinks. With the Venna programme (http://bioinfogp.cnb.csic.es), 146 of the 176 proteins were assigned exclusively to healthy dogs, and 373 of the 403 proteins were exclusively assigned to dogs with babesiosis; 30 proteins were common to both groups (Fig. 1). To further evaluate the 373 proteins found in only the dogs with babesiosis, the Panther programme (http://www.pantherdb.org) was used to isolate 21 proteins from the *Canis familiaris* species, which were used to form a collection of potential diagnostic and pathophysiological biomarkers for this disease (Table 3). Further analysis of these 21 proteins led to the isolation of 8 proteins associated with 10 metabolic pathways that were attributed to immune and inflammatory response development (Table 4). These results showed that the epithelial-mesenchymal transition (EMT) might play an important role in the mechanisms underlying pathological changes in renal tissues during the course of babesiosis, as indicated by the causal relationship network built by combining 5 of the 10 selected metabolic pathways and 4 of the 8 proteins for which the pathways were associated. These included cadherins, gonadotropin releasing hormone receptors, inflammatory responses mediated by chemokine and cytokine signalling pathways, integrins, and TGF-β pathways. These pathways were linked by interleukin (IL)-13, bone morphogenetic protein 7, α2(1) collagen, and FER tyrosine kinase.

**Table 1.**
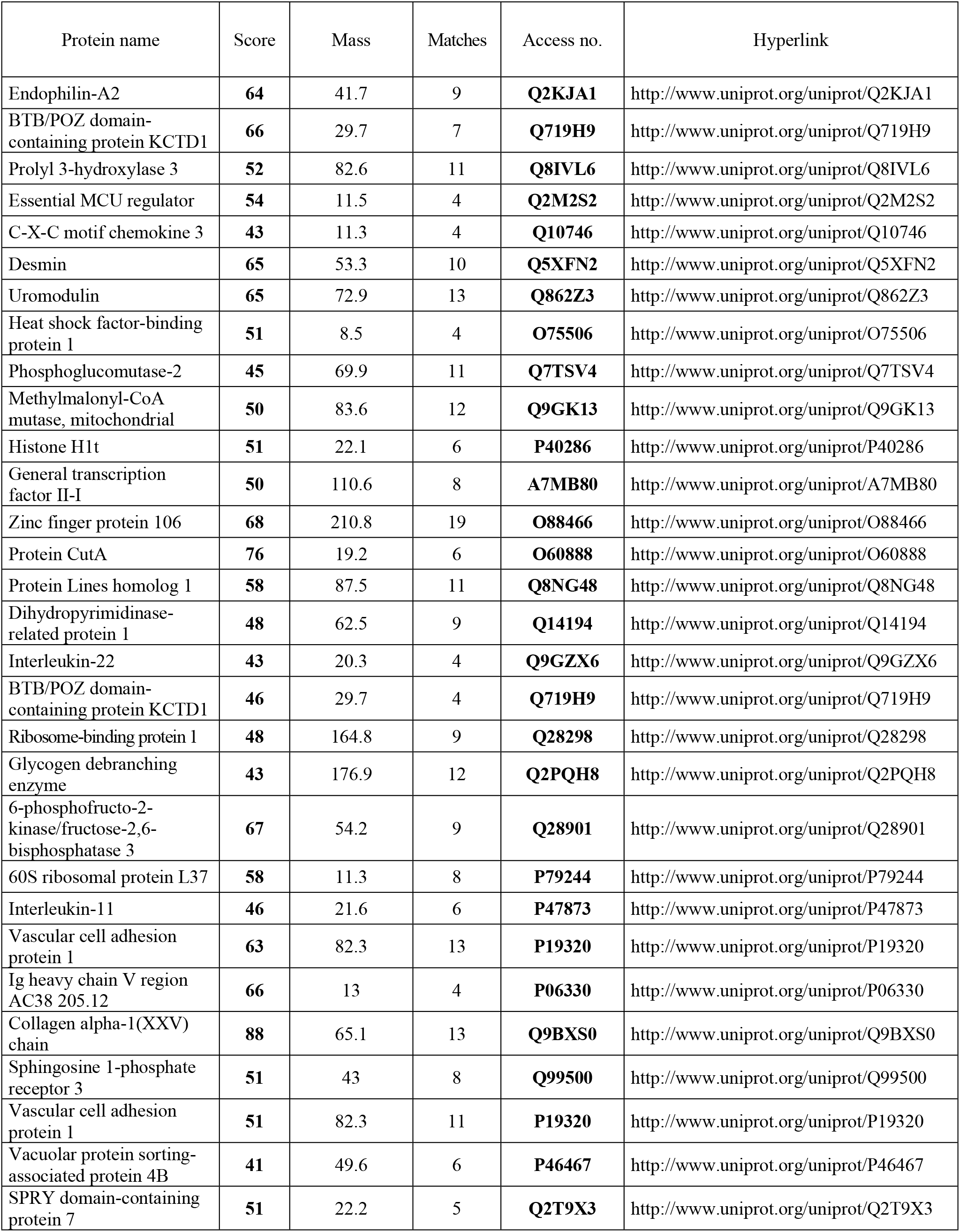
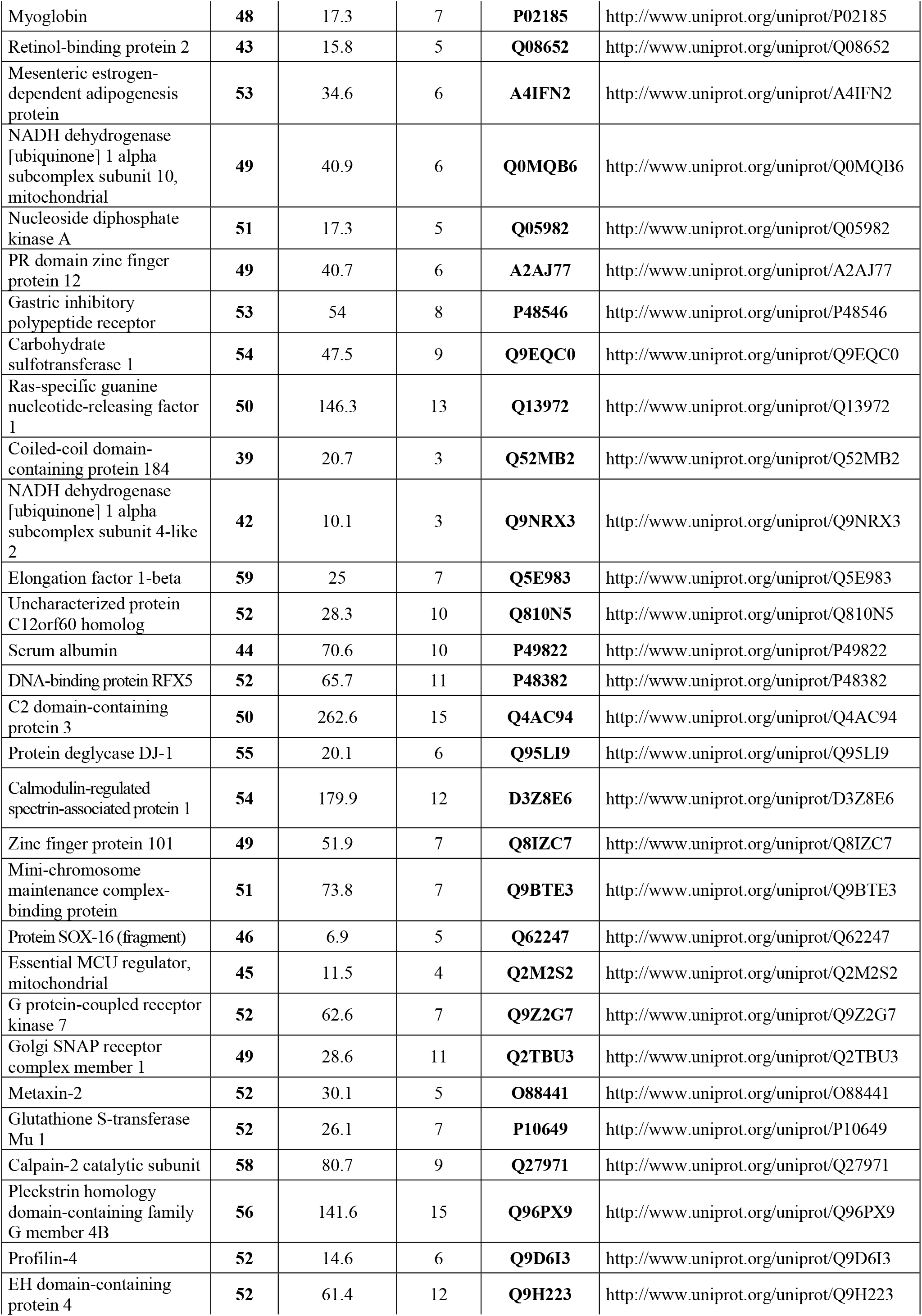
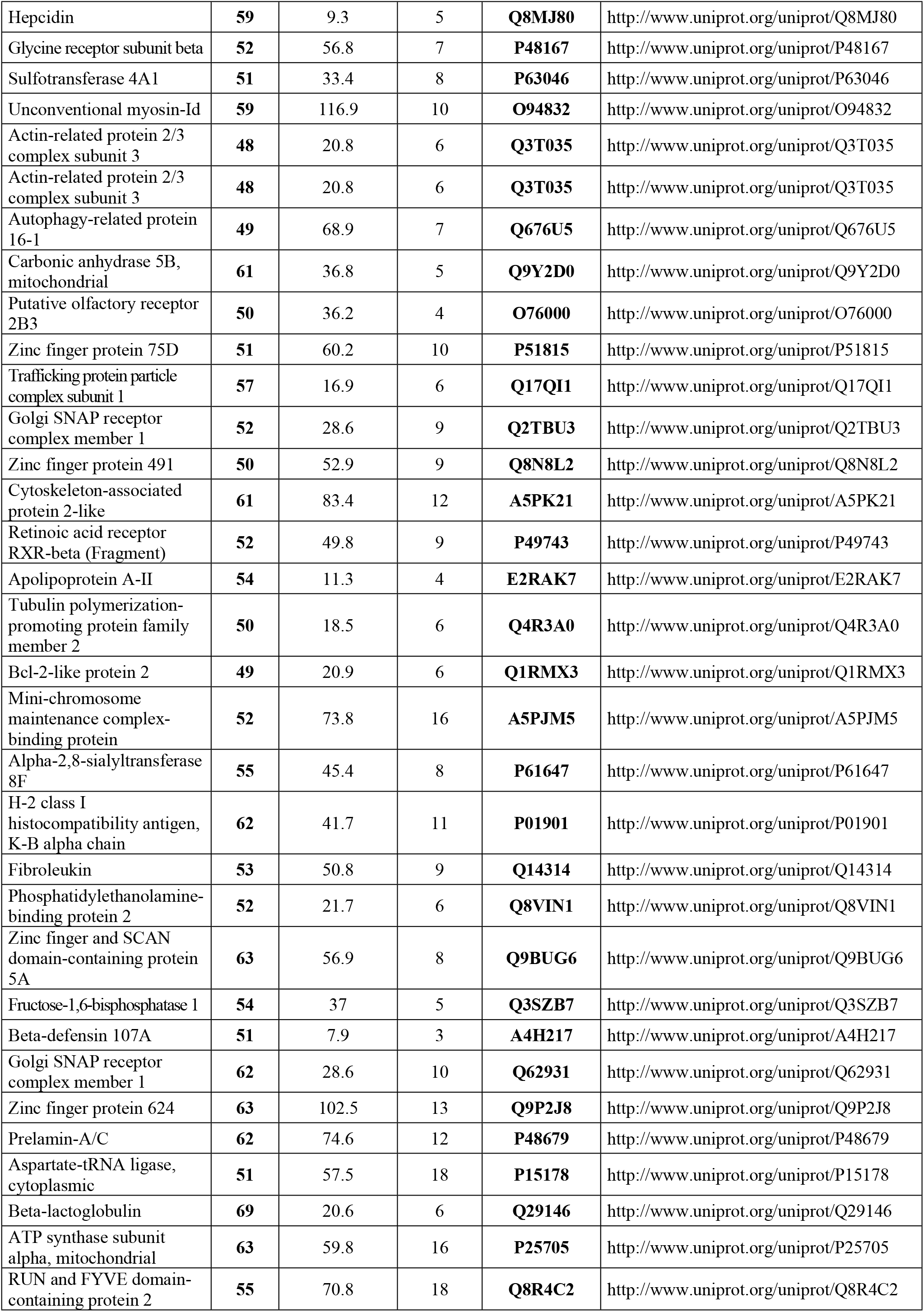
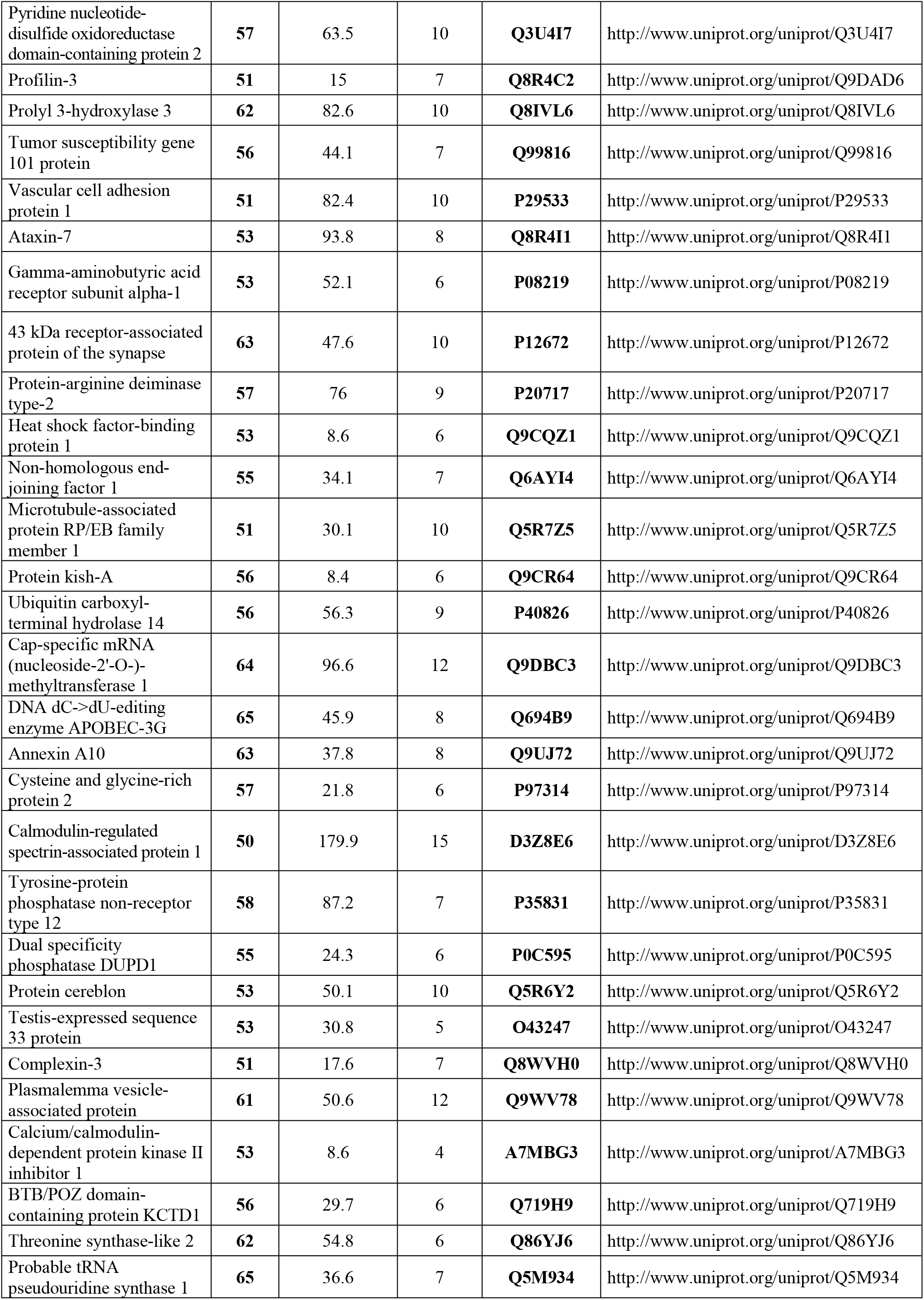
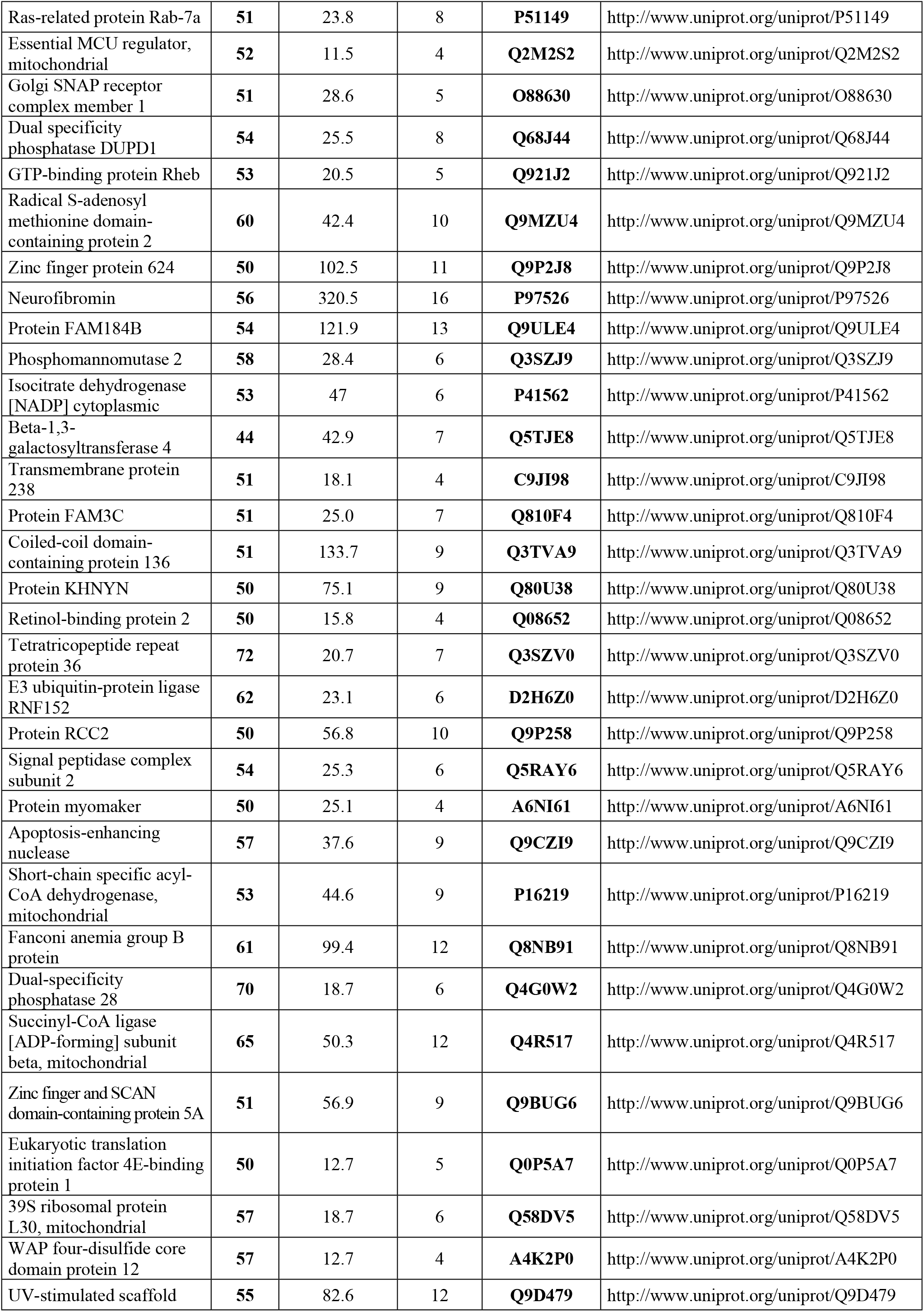
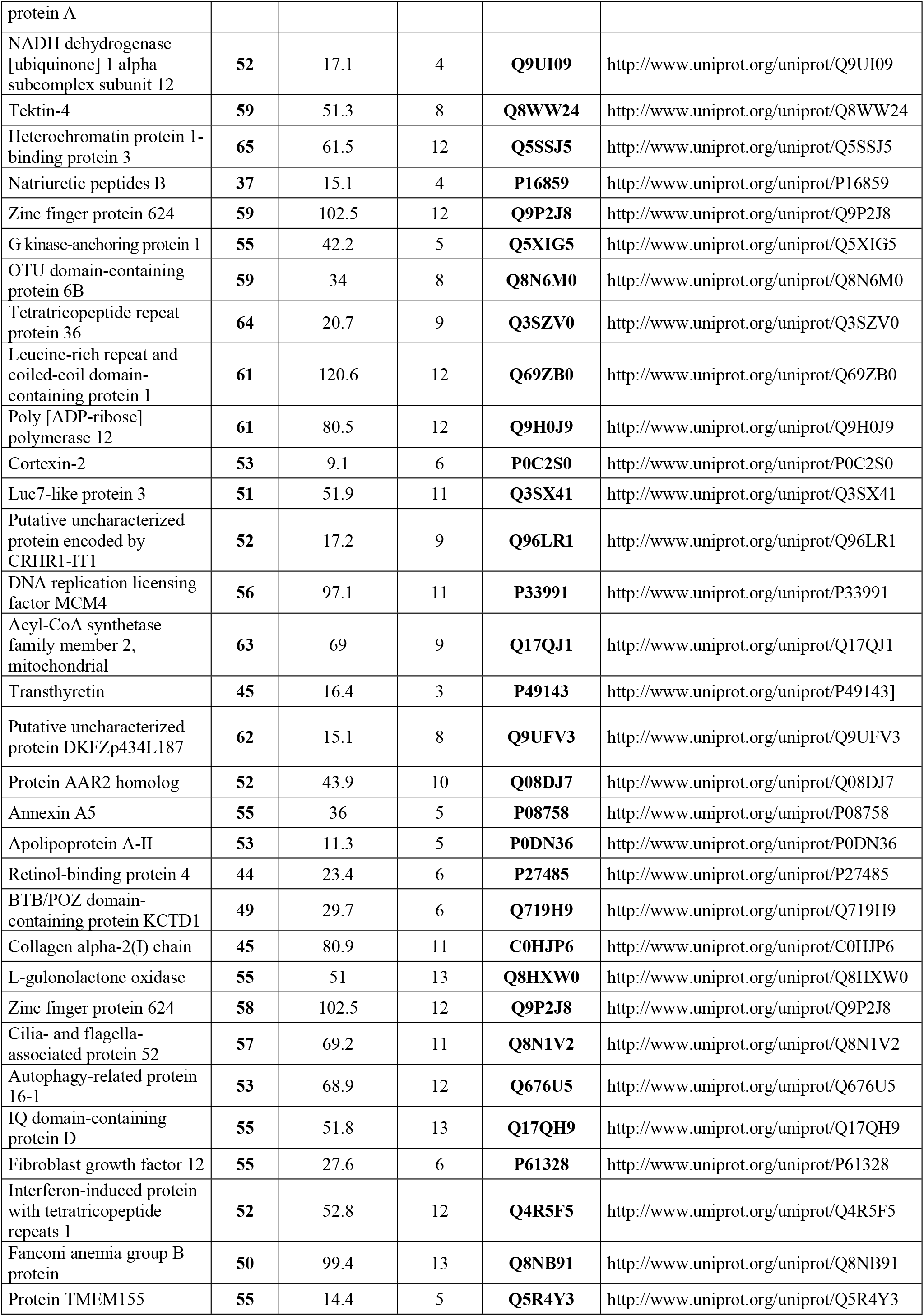
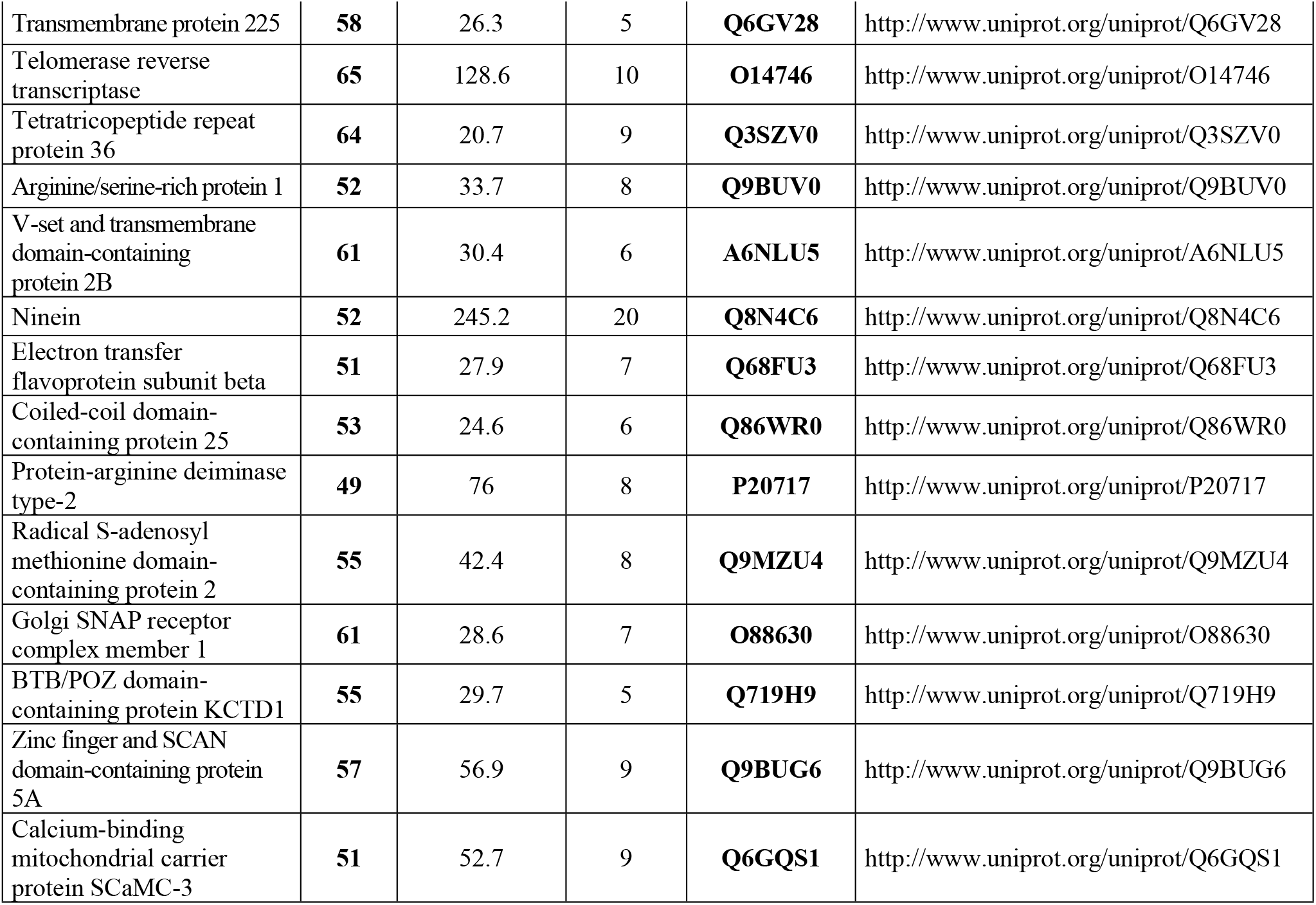
Peptides identified in the urine of healthy dogs.

**Table 2.**
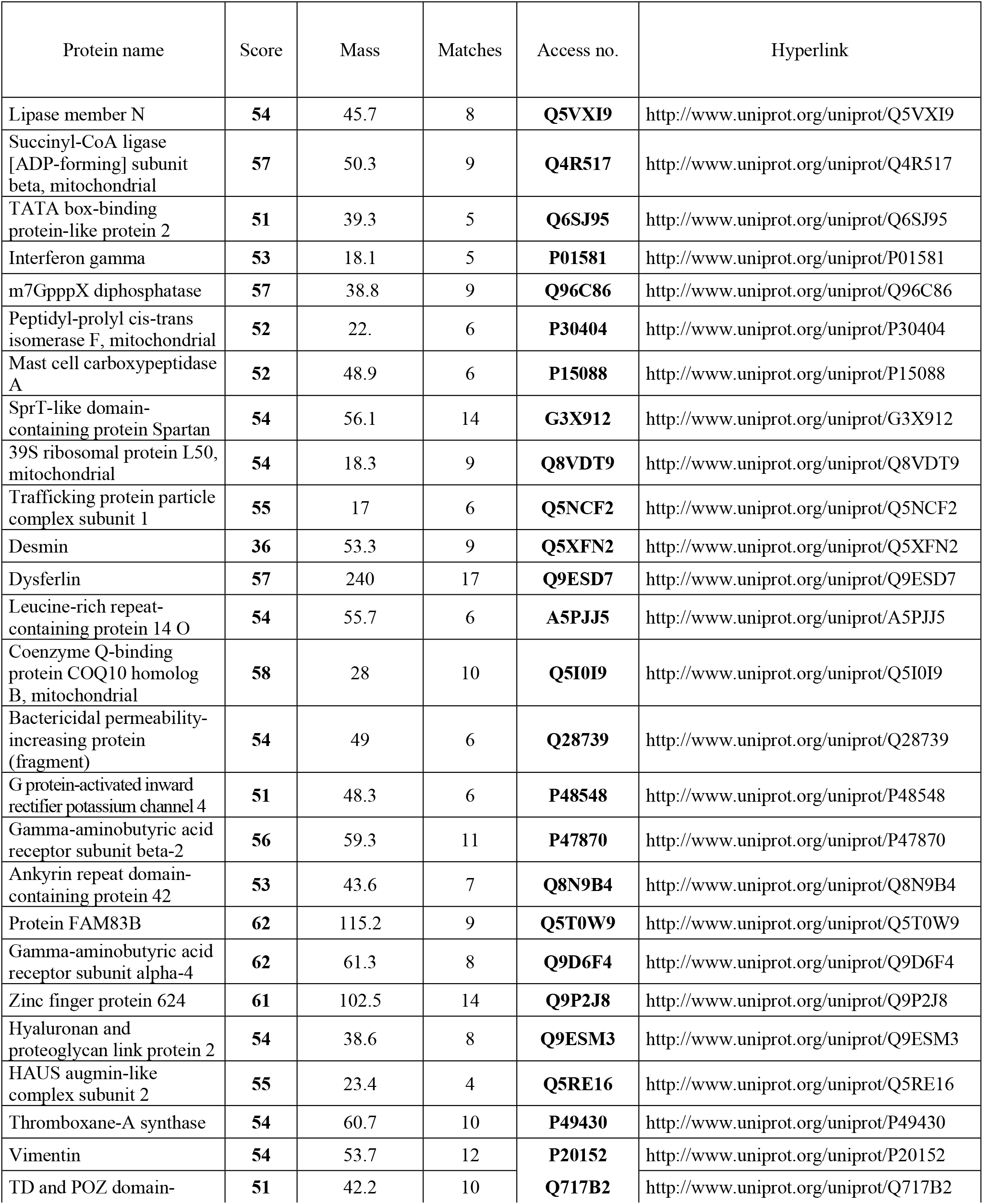
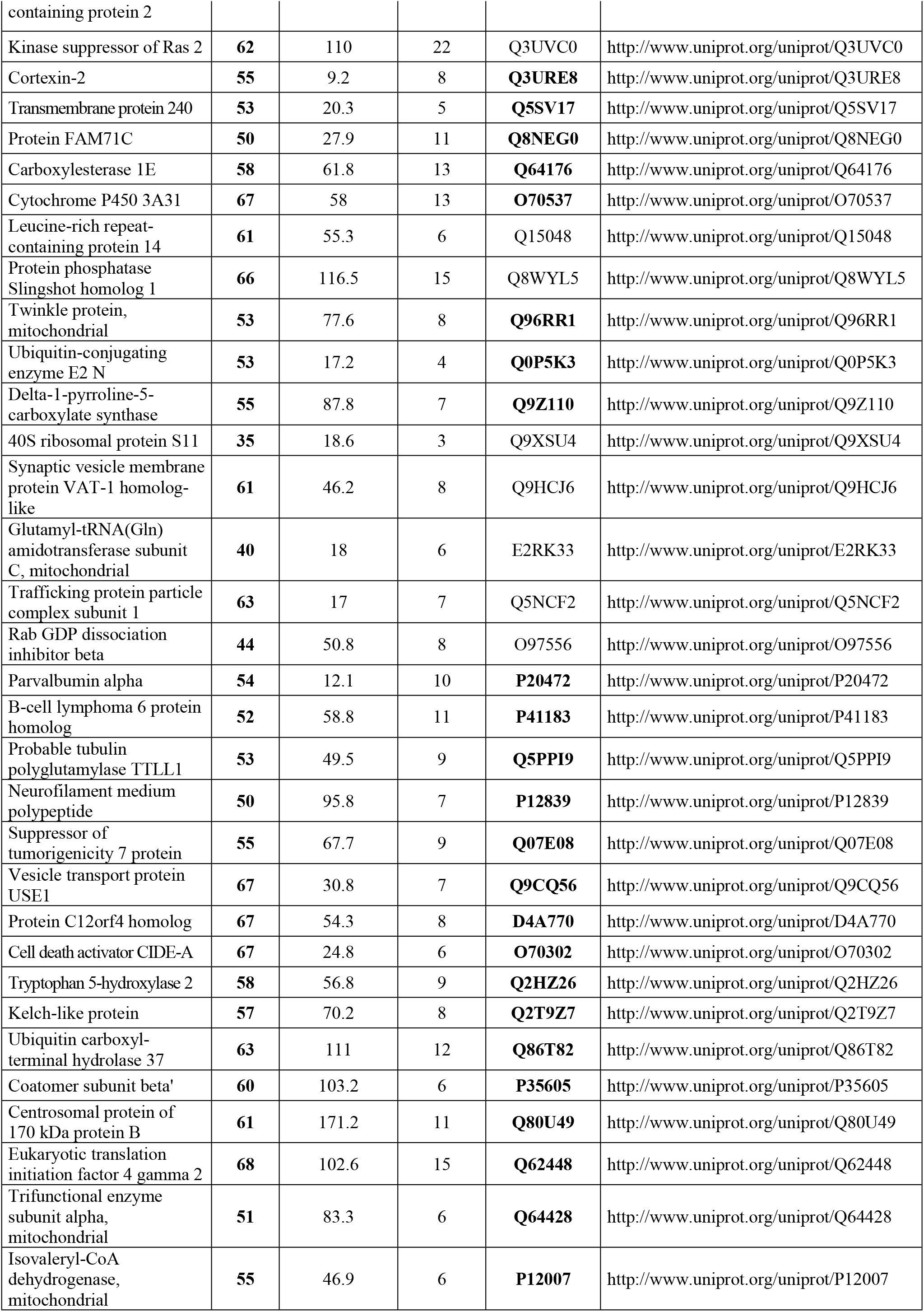
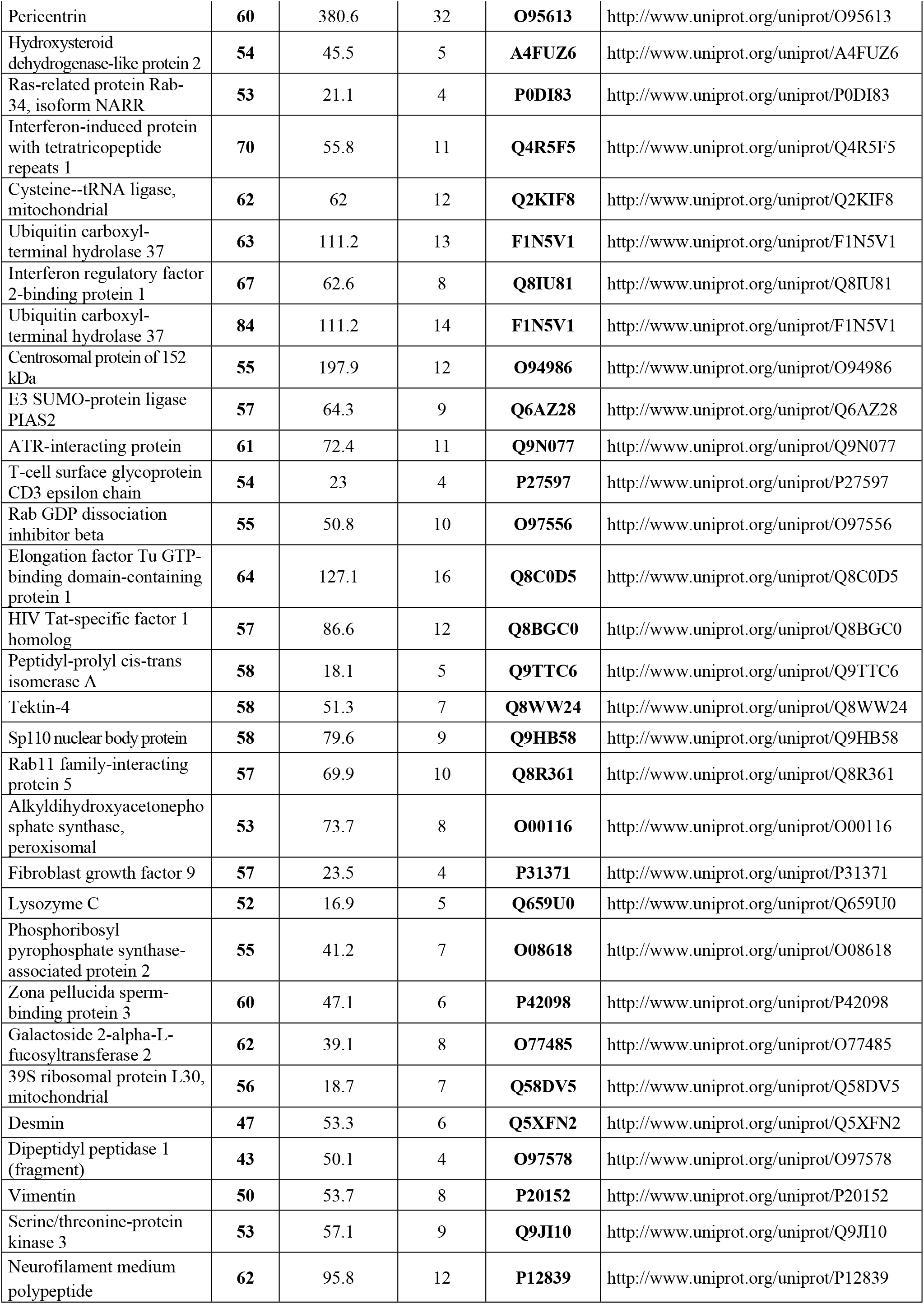
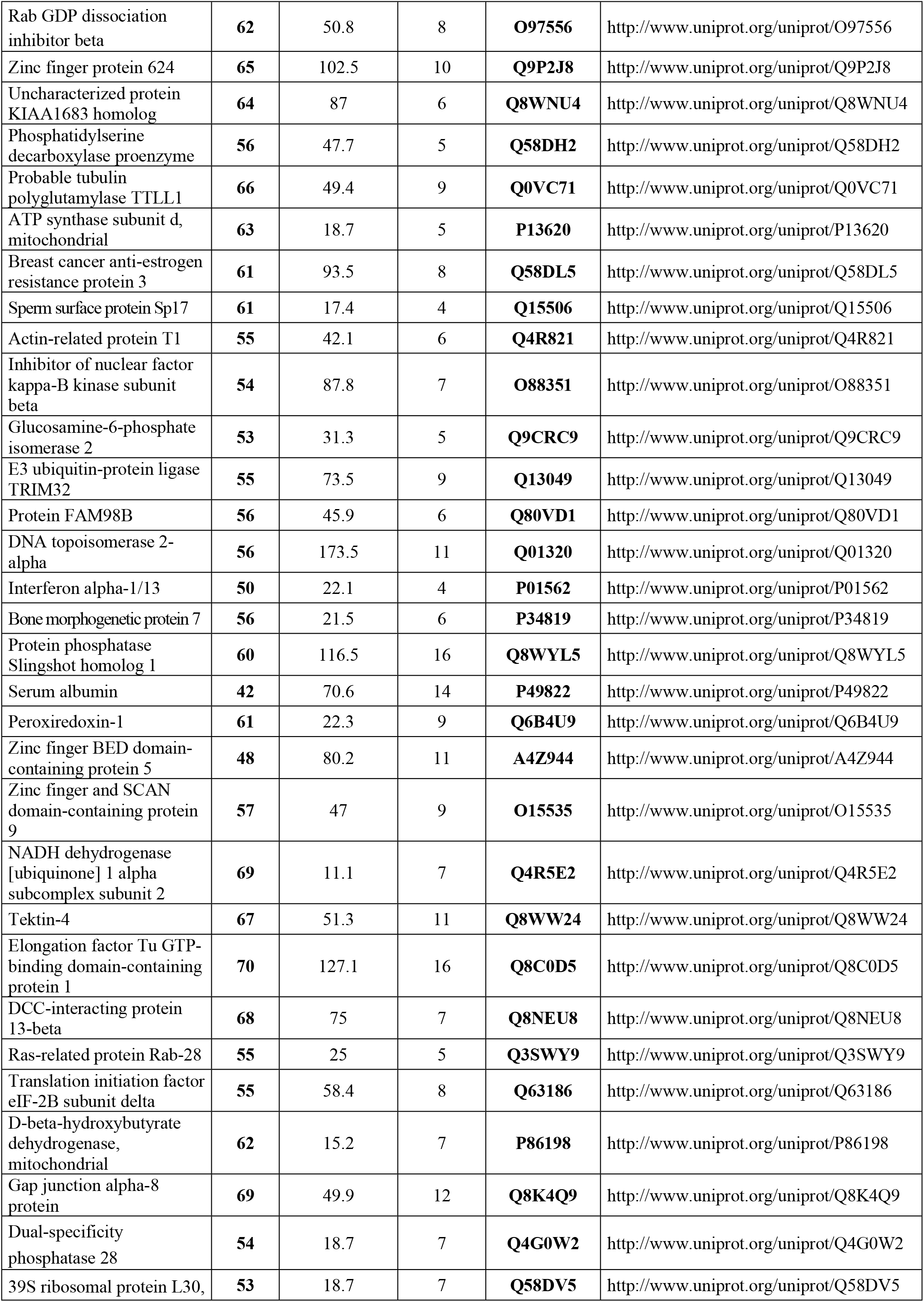
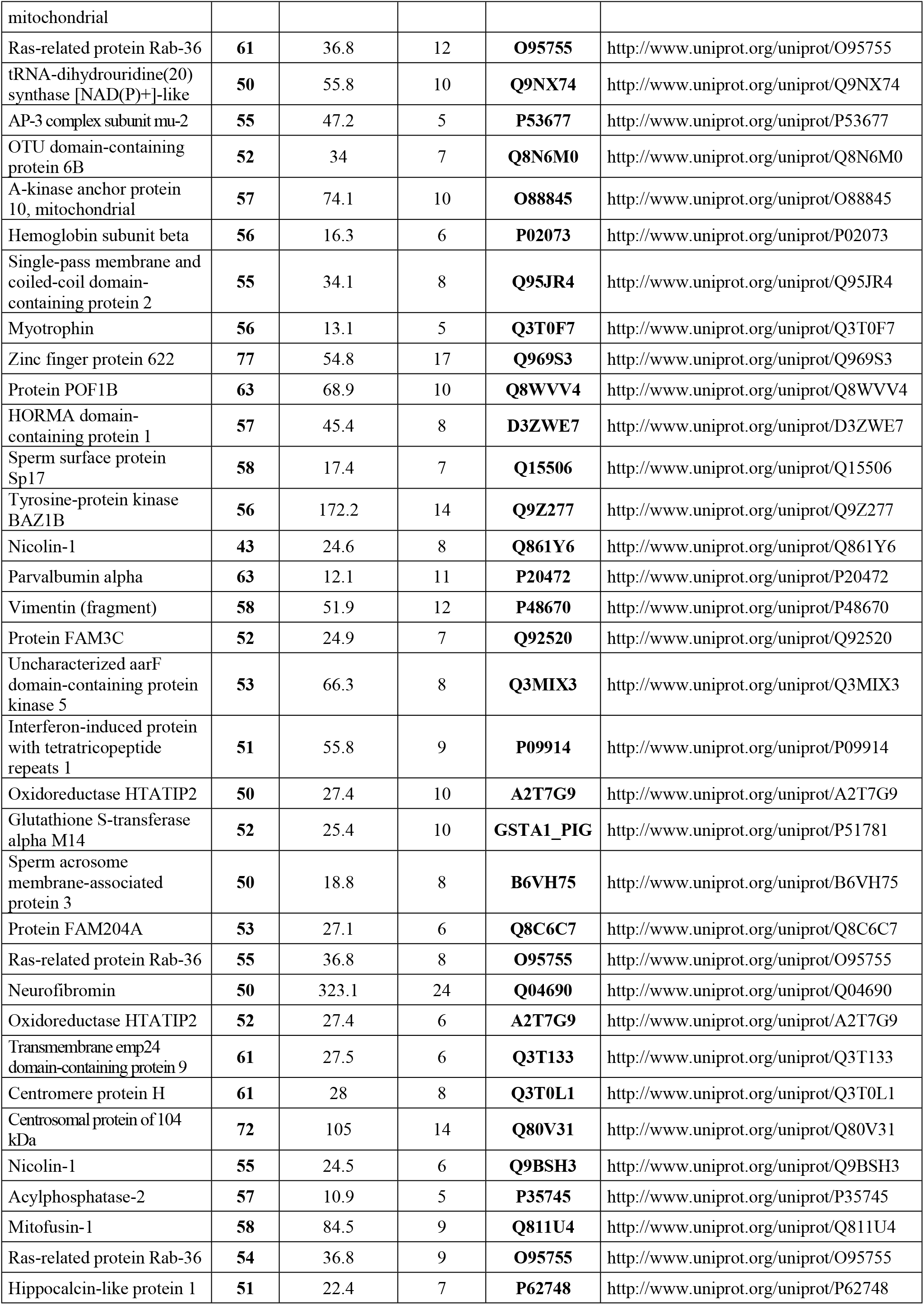
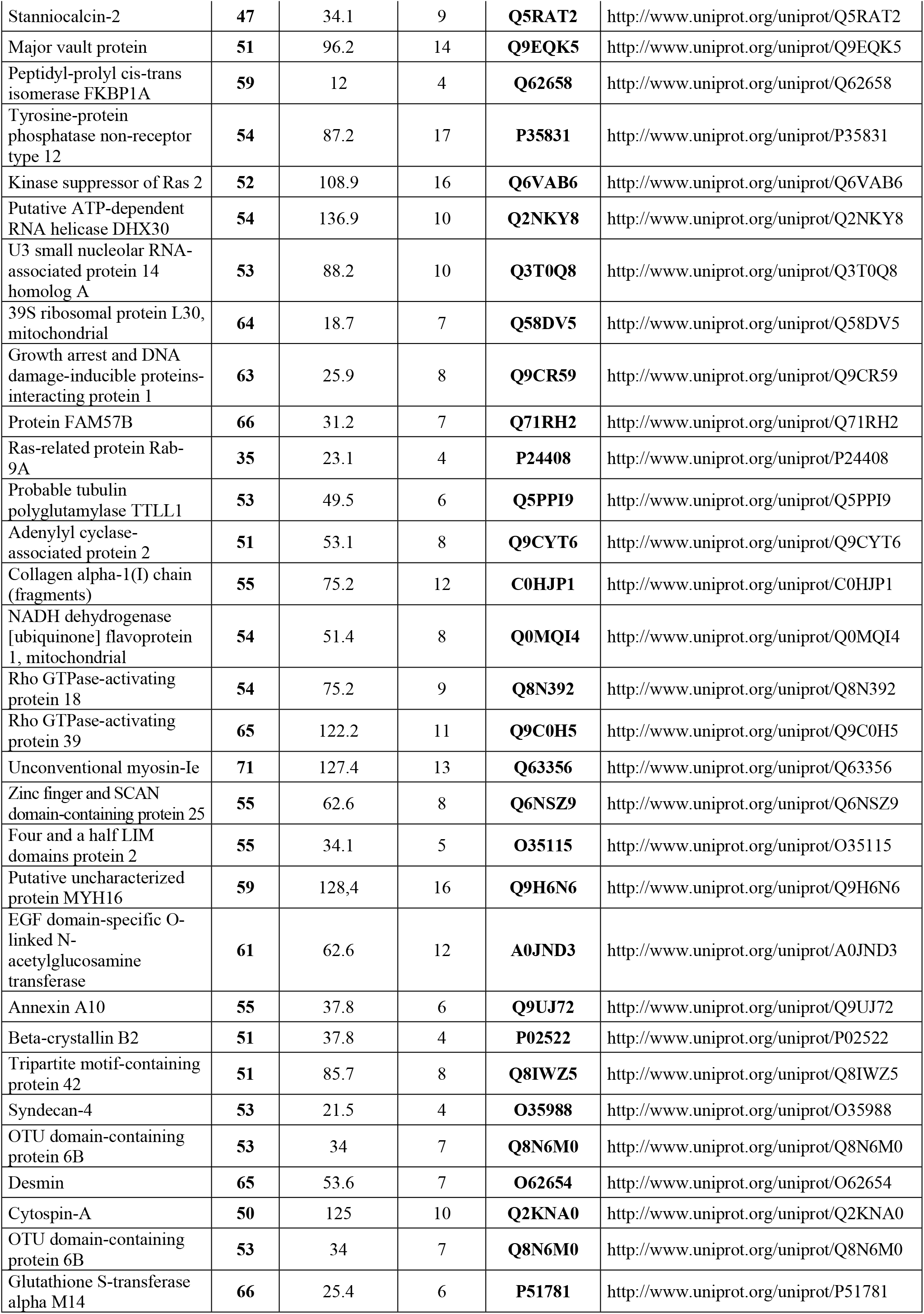
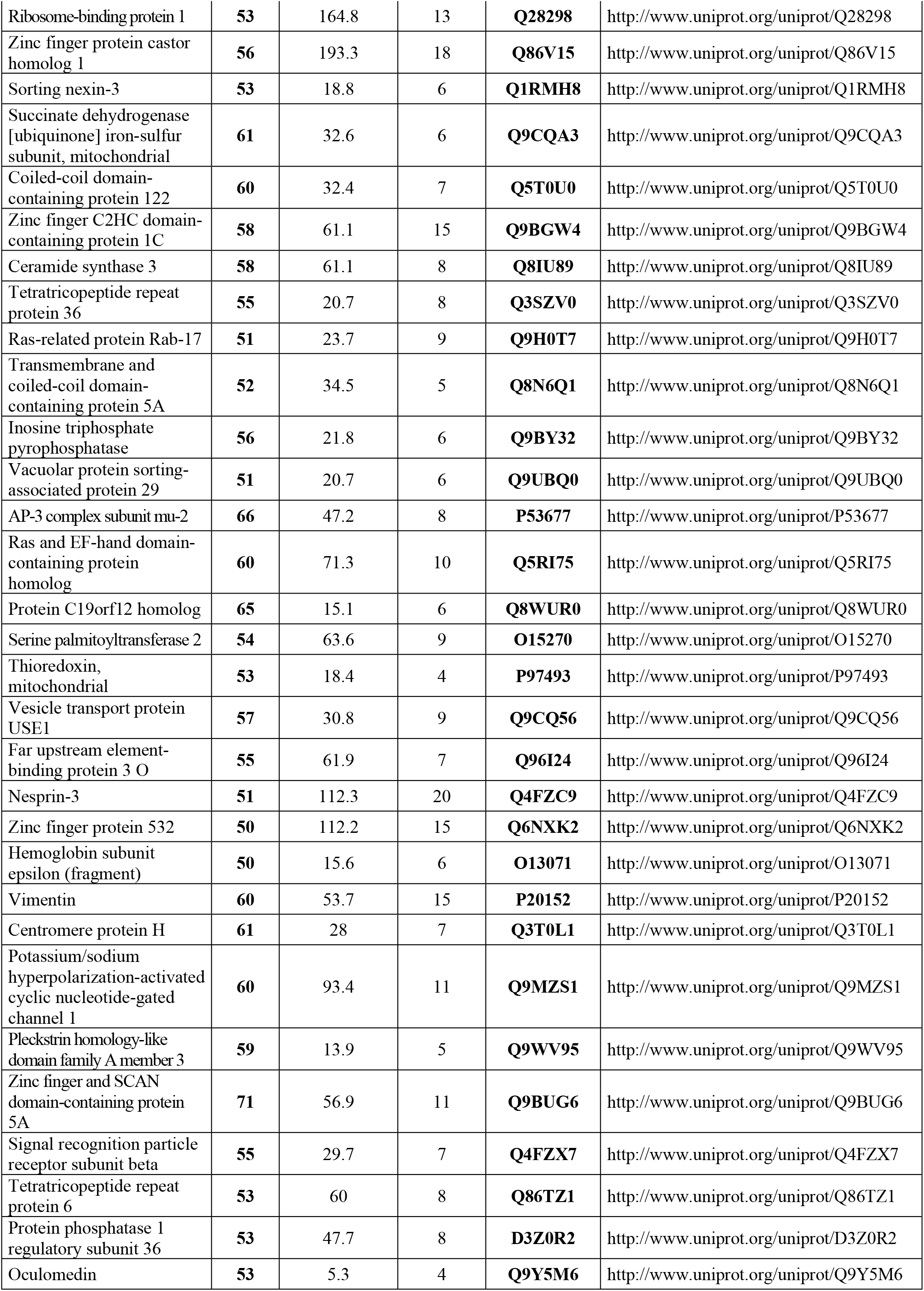
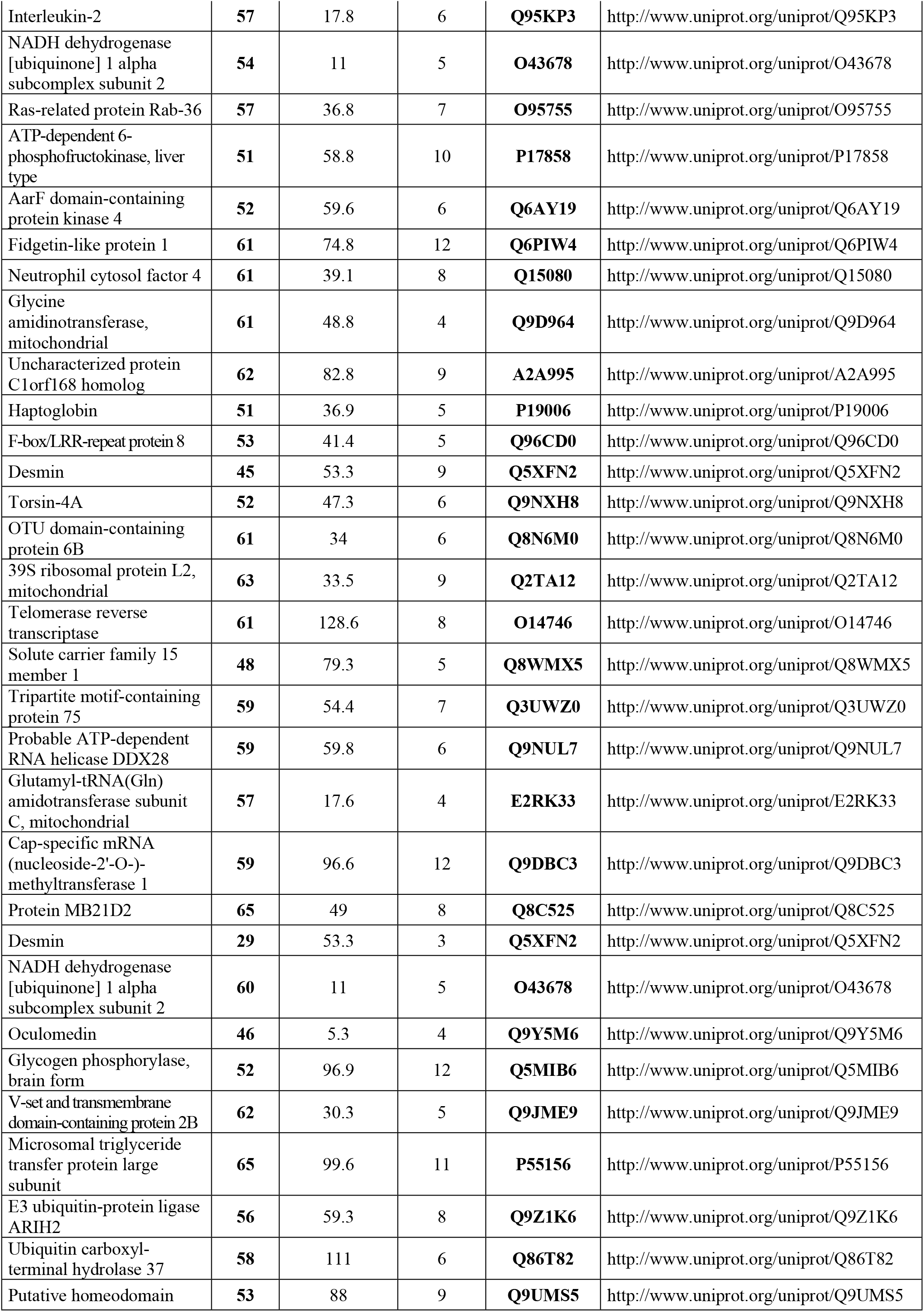
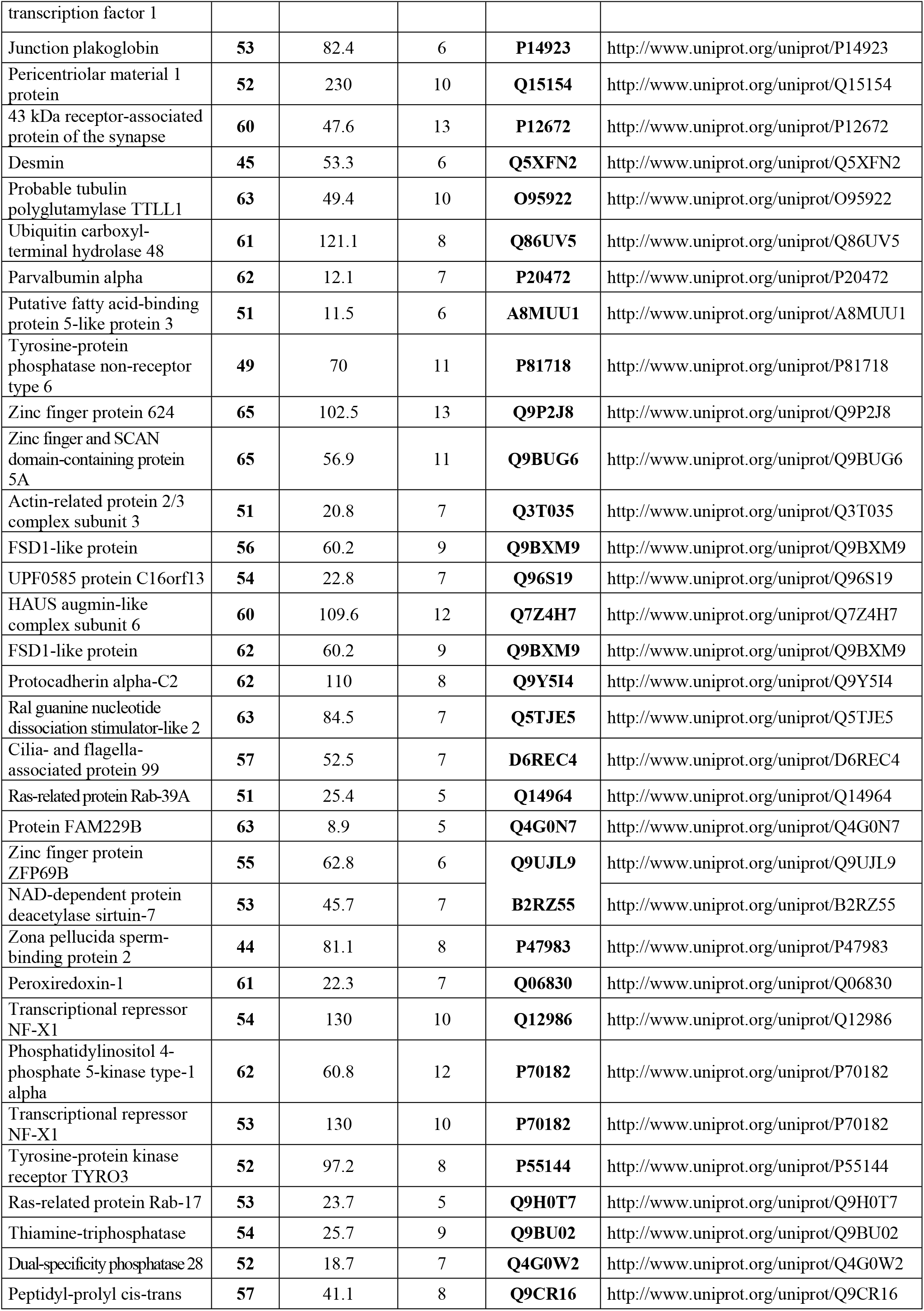
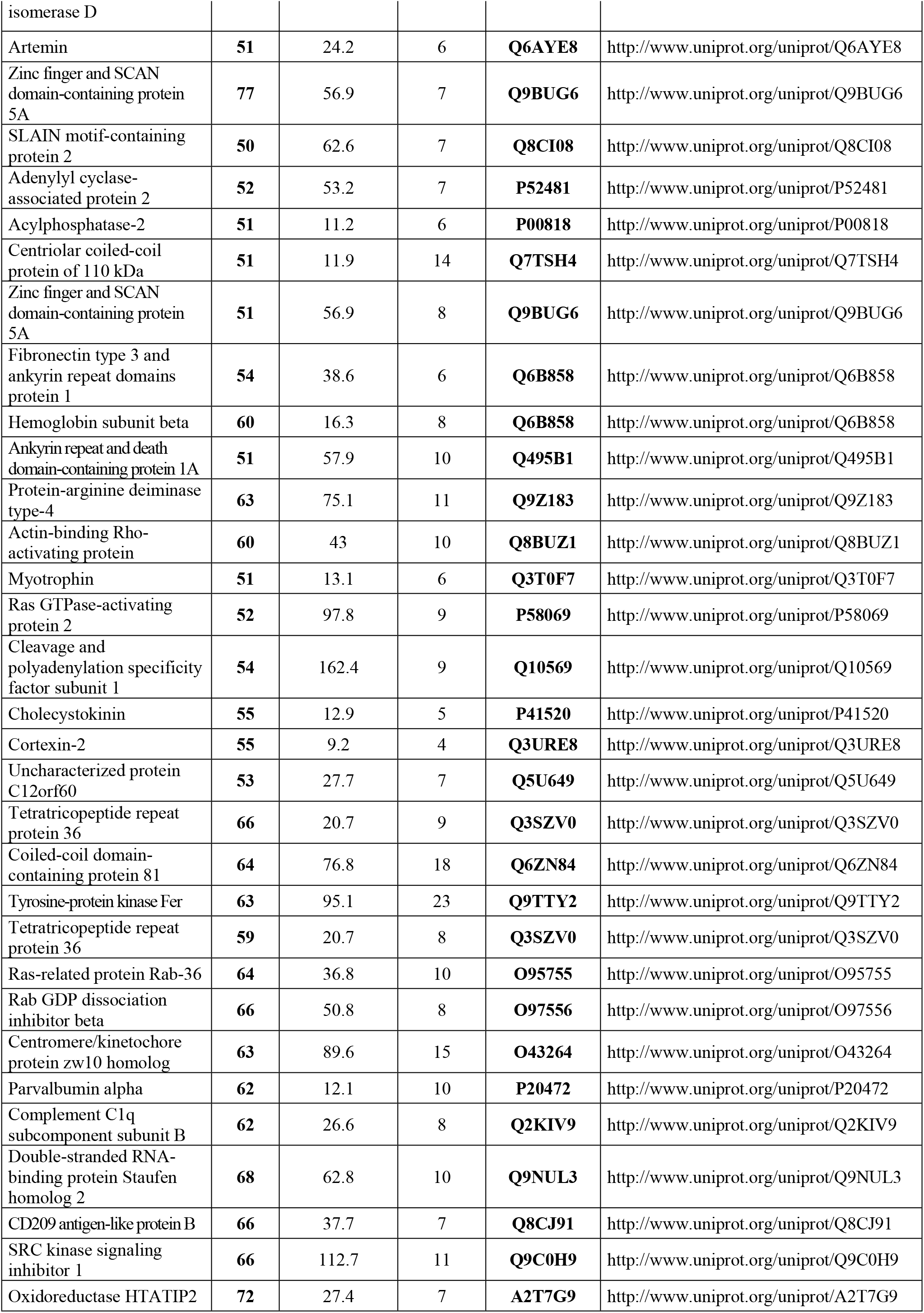
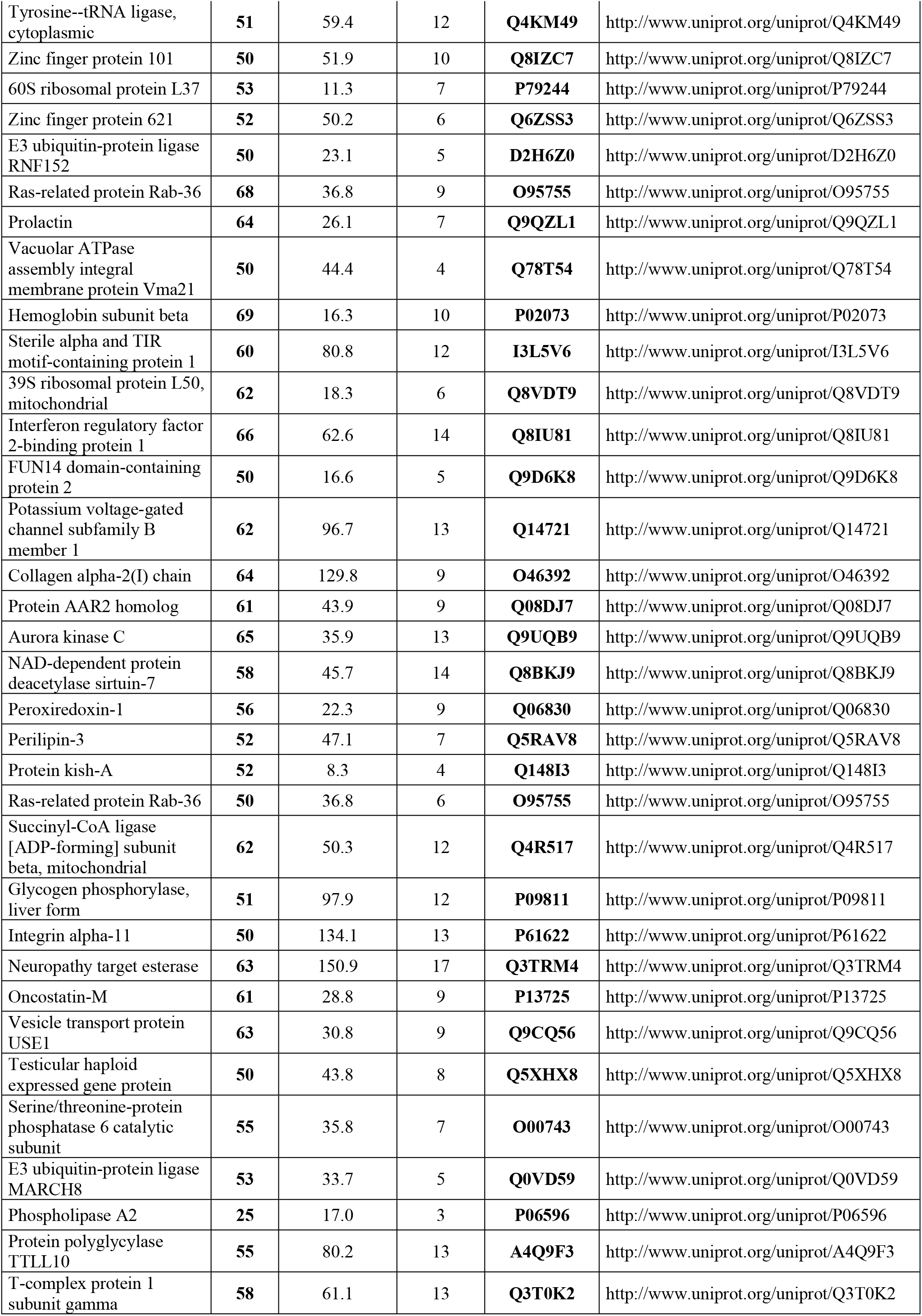
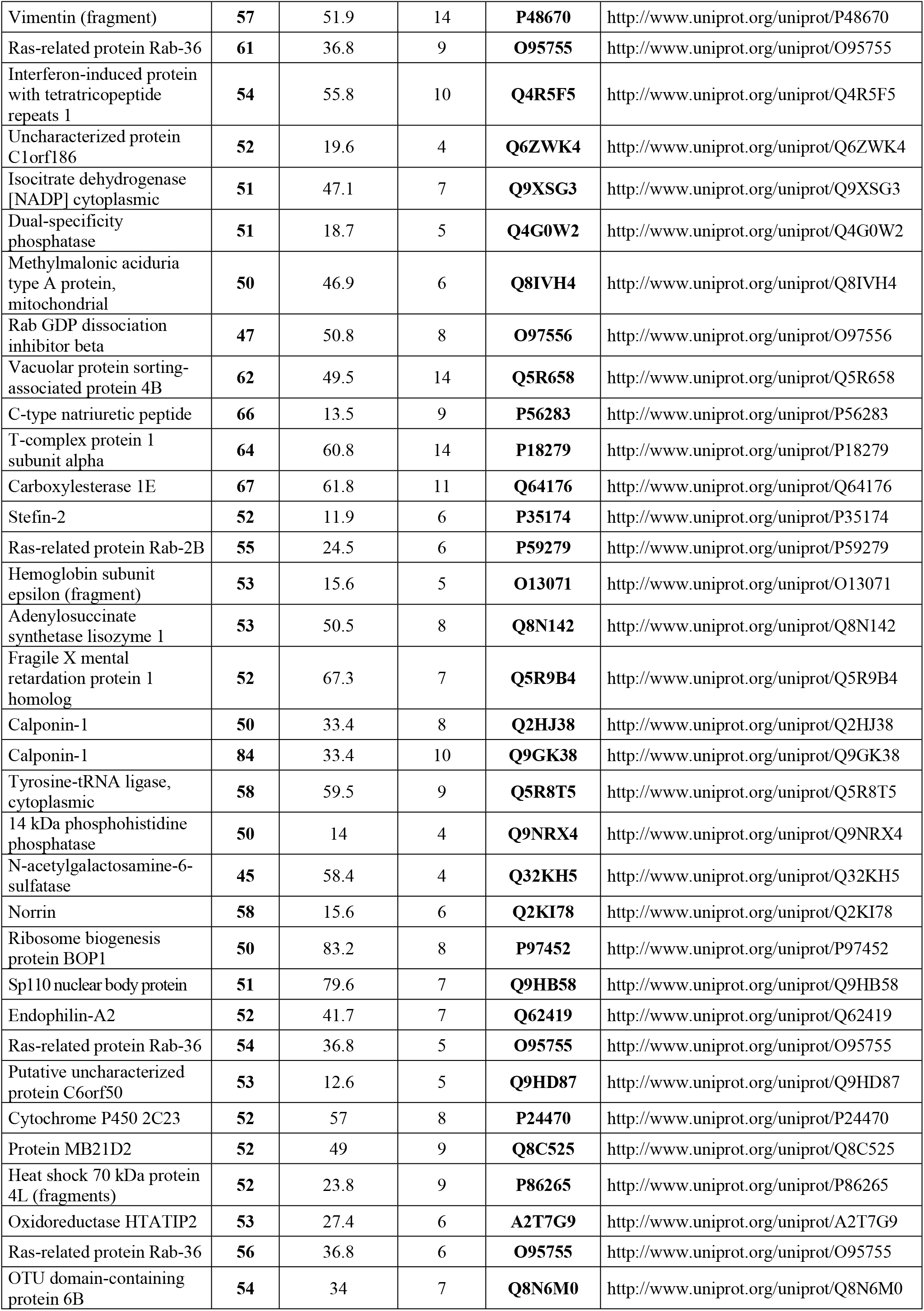
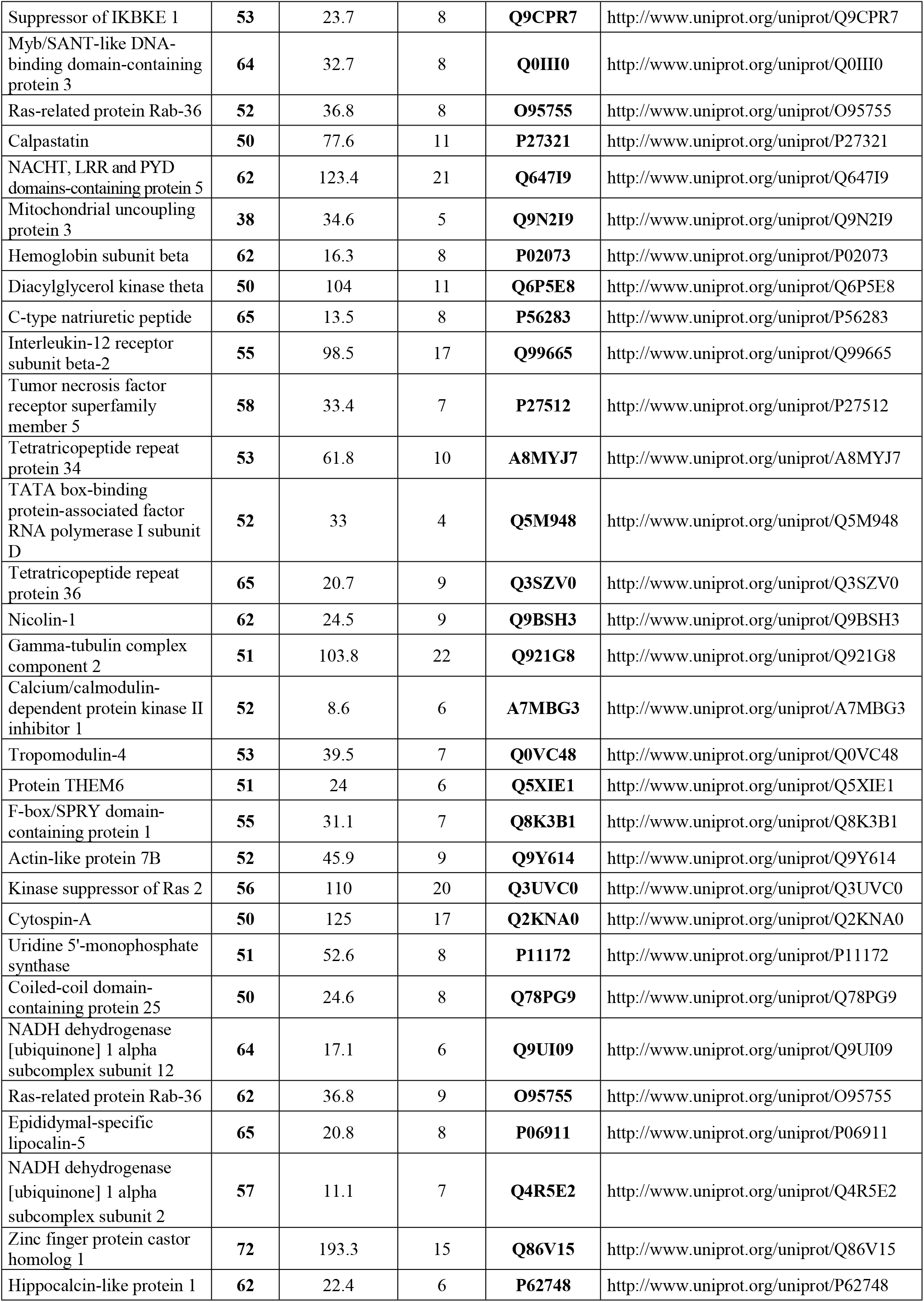
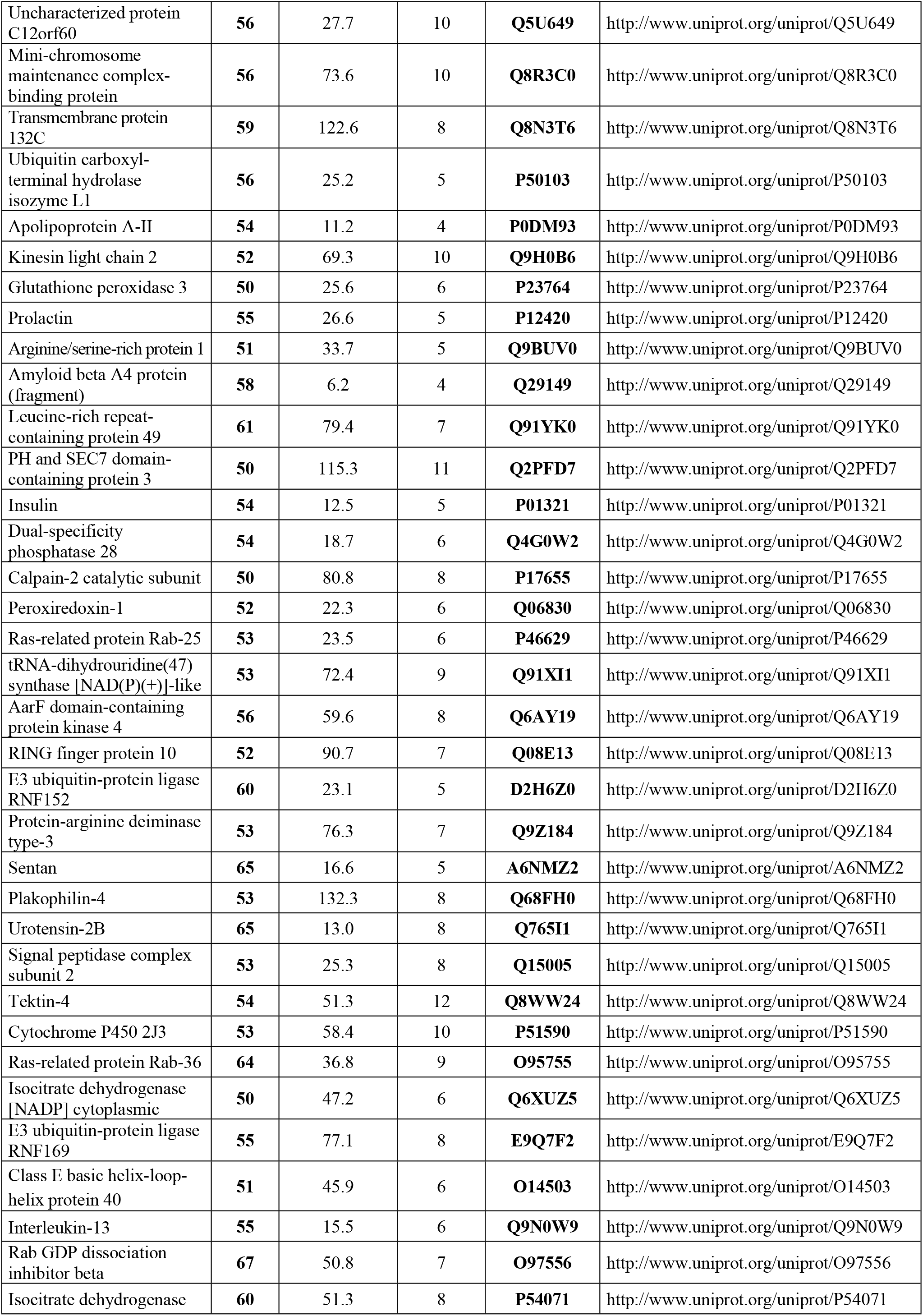
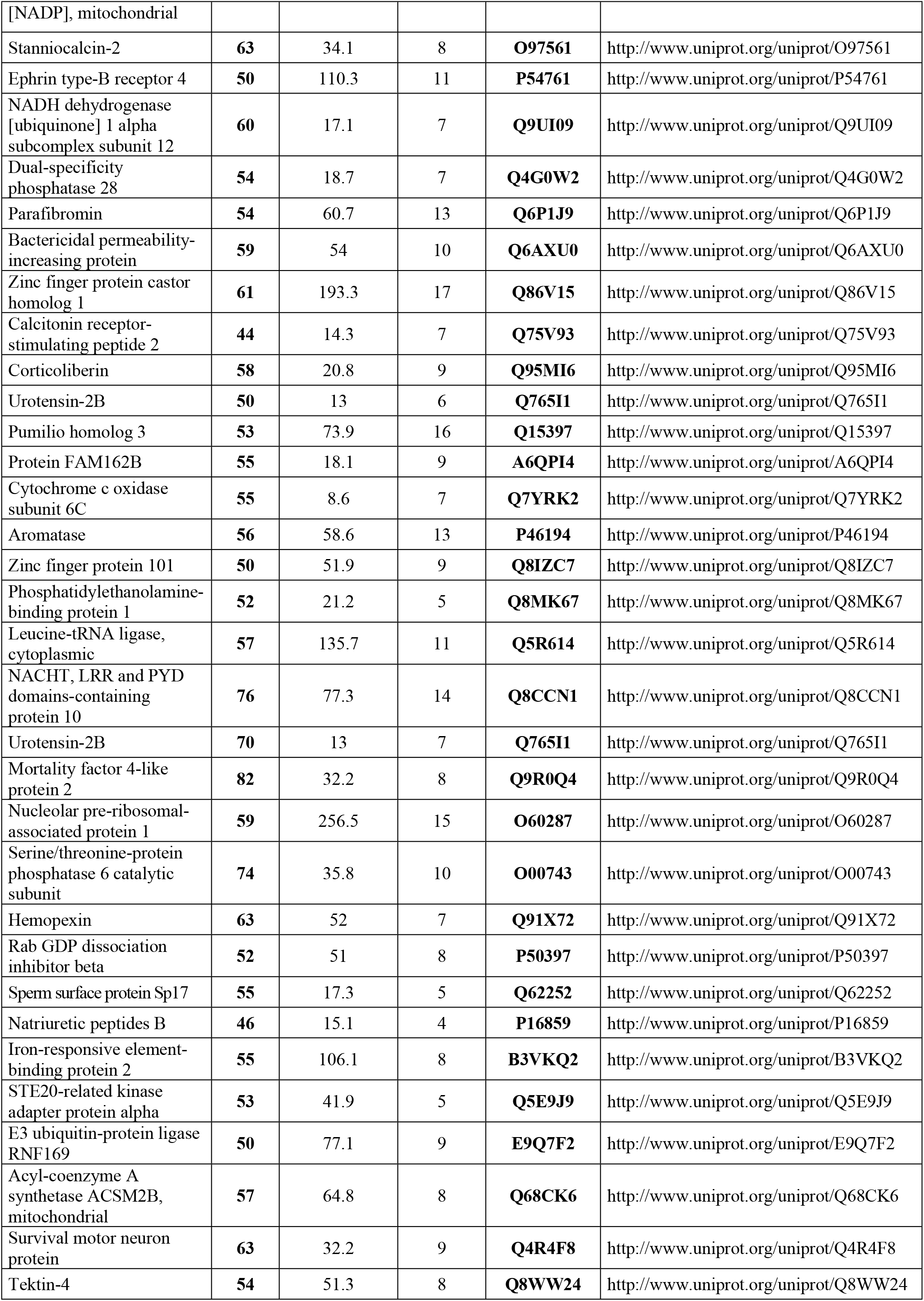
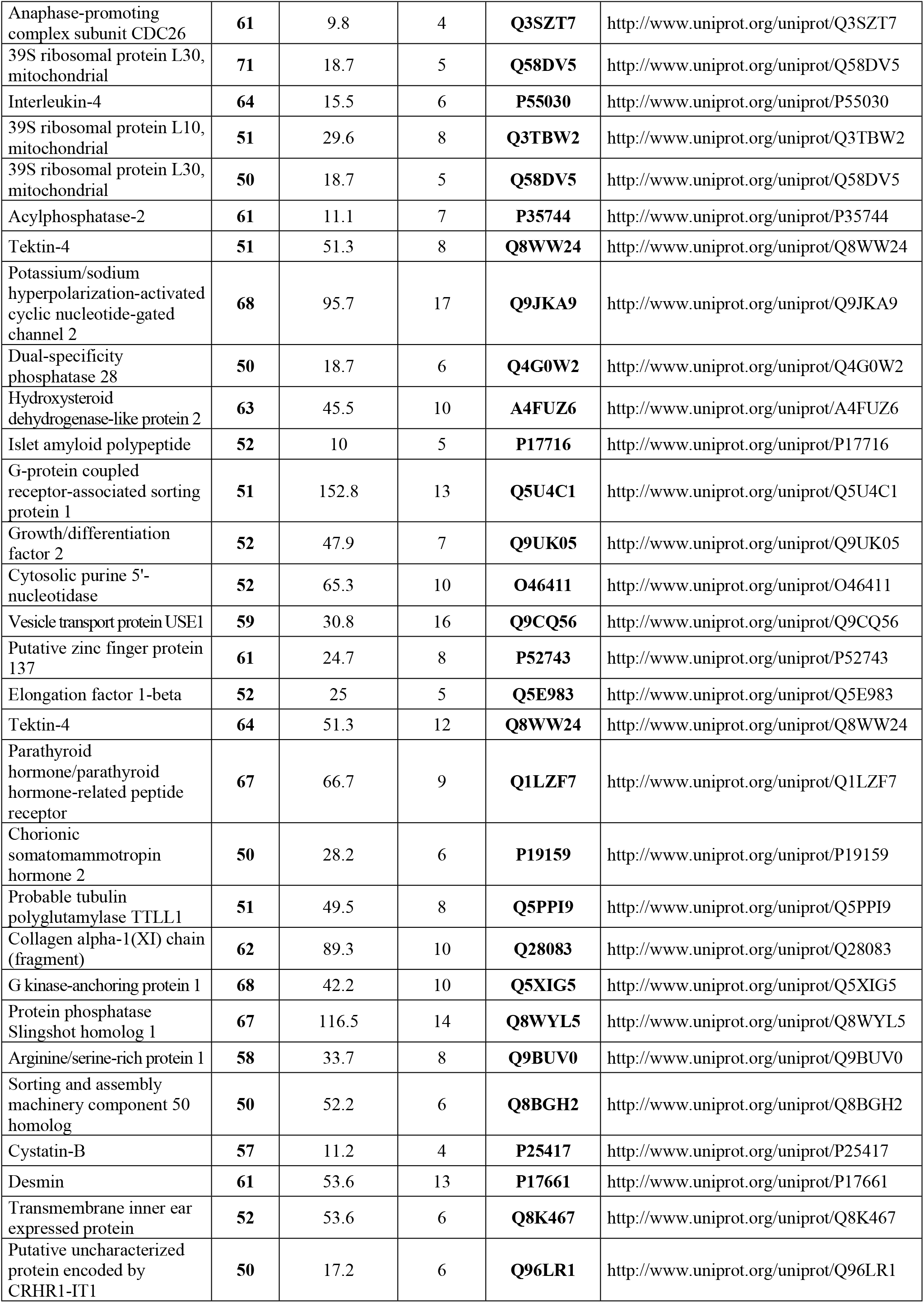
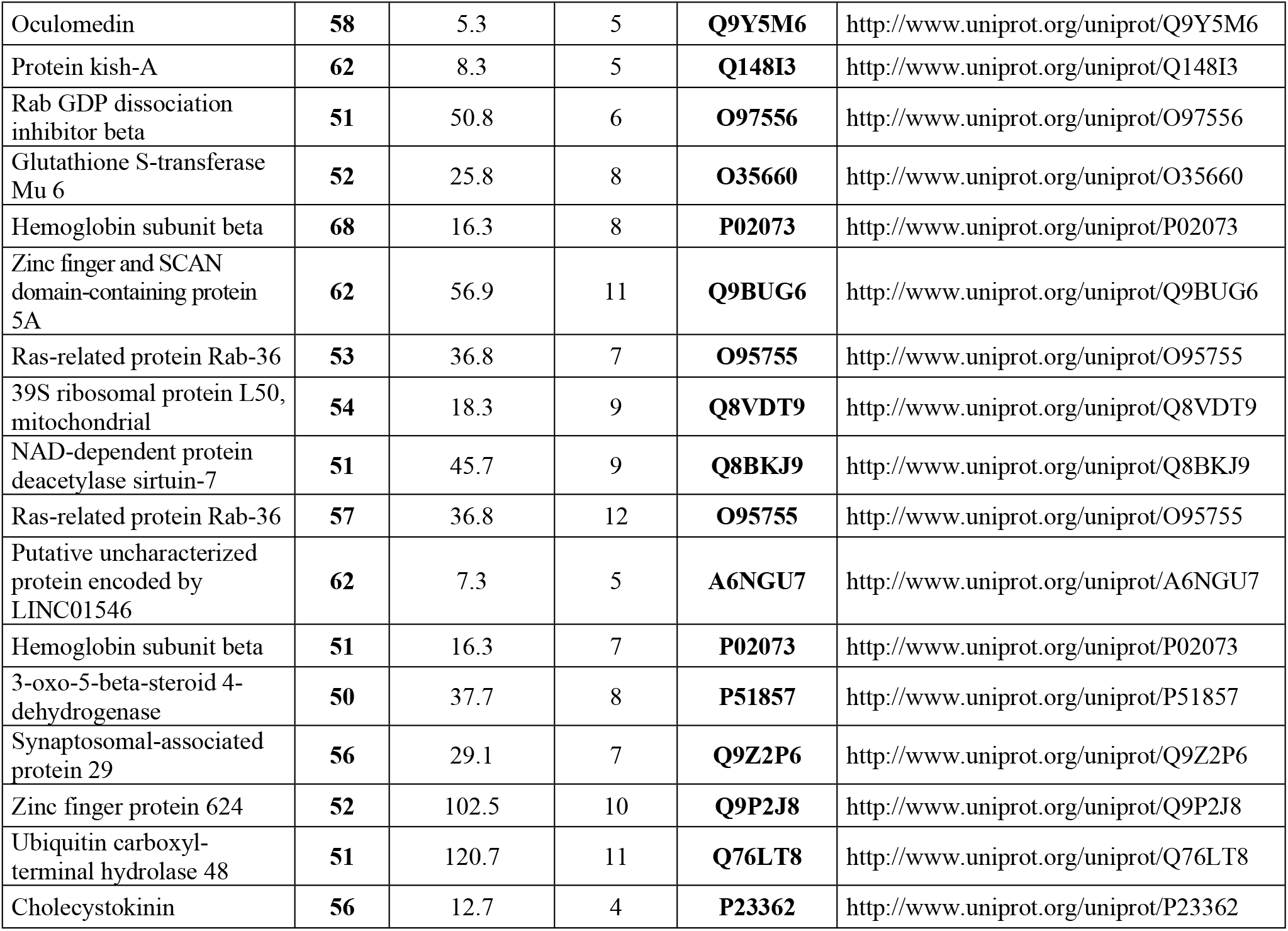
Peptides identified in the urine of dogs with babesiosis.

**Table 3.** List of *Canis familiaris* proteins identified in the urine of dogs with babesiosis by MALDI-TOF/TOF.

| Nr | Accession<sup>a</sup> | Protein name | GO molecular function |
| --- | --- | --- | --- |
| 1 | P06596 | Phospholipase A2 | phospholipase |
| 2 | A4Z944 | Zinc finger BED domain-containing protein 5 | transcription factor |
| 3 | O46392 | Collagen alpha-2(I) chain | extracellular matrix structural constituent |
| 4 | Q9XSU4 | 40S ribosomal protein S11 | structural constituent of ribosome |
| 5 | O97556 | Rab GDP dissociation inhibitor beta | G-protein modulator acyltransferase |
| 6 | Q9TTY2 | Tyrosine-protein kinase Fer | tyrosine kinase activity |
| 7 | P19006 | Haptoglobin | hemoglobin binding |
| 8 | Q75V93 | Calcitonin receptor-stimulating peptide 2 | peptide hormone |
| 9 | Q2KNA0 | Cytospin-A | structural component |
| 10 | P01321 | Insulin | hormone |
| 12 | Q9N2I9 | Mitochondrial uncoupling protein 3 | oxidative phosphorylation |
| 11 | E2RK33 | Glutamyl-tRNA(Gln) amidotransferase subunit C, mitochondrial | ligase |
| 12 | Q32KH5 | N-acetylgalactosamine-6-sulfatase | hydrolase |
| 13 | P27597 | T-cell surface glycoprotein CD3 epsilon chain | transmembrane signalling receptor |
| 14 | P24408 | Ras-related protein Rab-9A | GTPase |
| 15 | P34819 | Bone morphogenetic protein 7 (fragment) | growth factor |
| 16 | Q5TJE5 | Ral guanine nucleotide dissociation stimulator-like 2 | guanyl-nucleotide exchange factor |
| 17 | Q9N0W9 | Interleukin-13 | cytokine |
| 18 | Q8WMX5 | Solute carrier family 15 member 1 | transporter |
| 19 | O97578 | Dipeptidyl peptidase 1 (fragment) | endopeptidase |
| 20 | Q861Y6 | Nicolin-1 | structural component |
| 21 | P17716 | Islet amyloid polypeptide | hormone |

a. in UniProt database (http://www.uniprot.org)

**Table 4.**
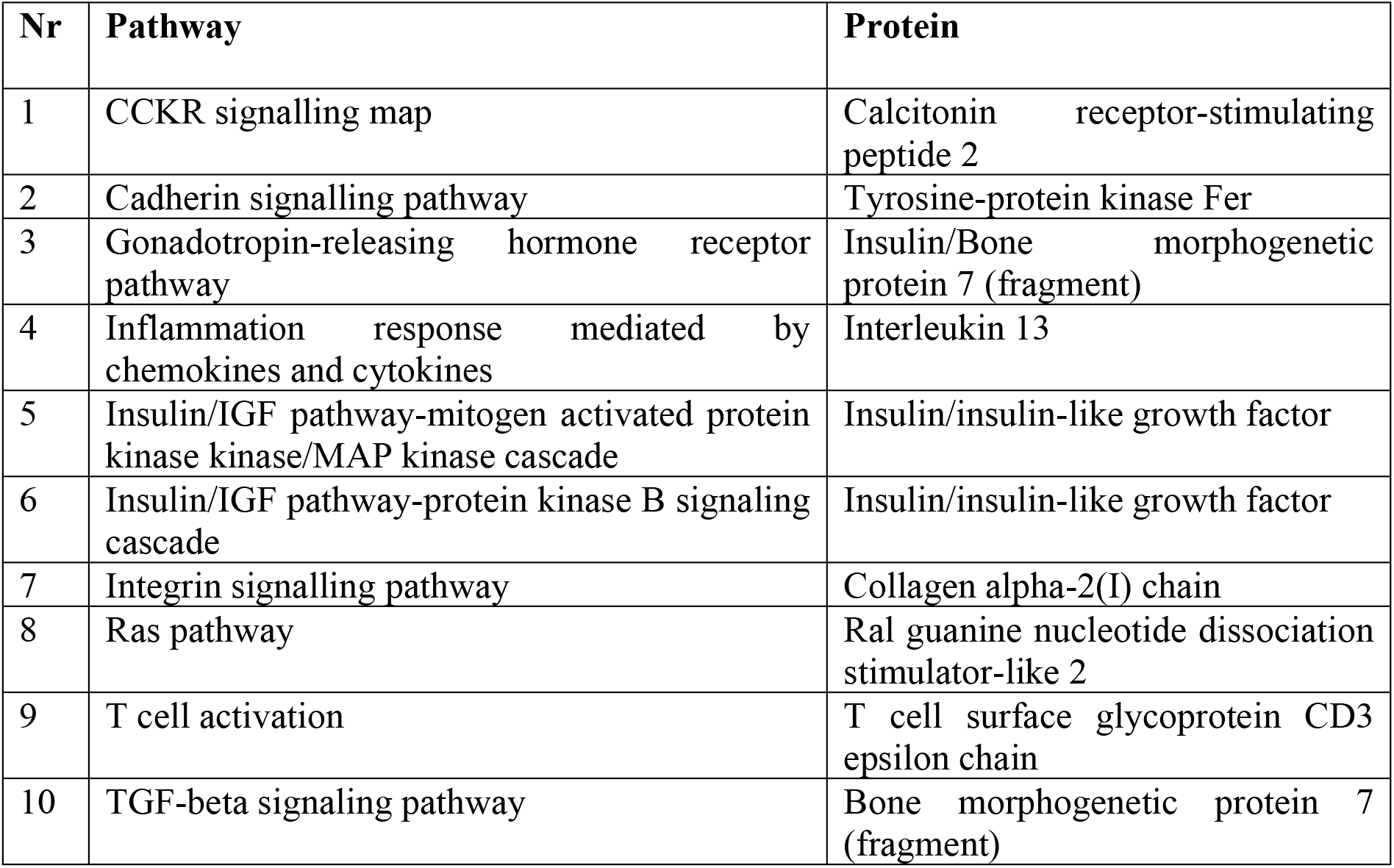
List of 10 metabolic pathways and 8 associated proteins in the urine of dogs with babesiosis.

| Nr | Pathway | Protein |
| --- | --- | --- |
| 1 | CCKR signalling map | Calcitonin receptor-stimulating peptide 2 |
| 2 | Cadherin signalling pathway | Tyrosine-protein kinase Fer |
| 3 | Gonadotropin-releasing hormone receptor pathway | Insulin/Bone morphogenetic protein 7 (fragment) |
| 4 | Inflammation response mediated by chemokines and cytokines | Interleukin 13 |
| 5 | Insulin/IGF pathway-mitogen activated protein kinase kinase/MAP kinase cascade | Insulin/insulin-like growth factor |
| 6 | Insulin/IGF pathway-protein kinase B signaling cascade | Insulin/insulin-like growth factor |
| 7 | Integrin signalling pathway | Collagen alpha-2(I) chain |
| 8 | Ras pathway | Ral guanine nucleotide dissociation stimulator-like 2 |
| 9 | T cell activation | T cell surface glycoprotein CD3 epsilon chain |
| 10 | TGF-beta signaling pathway | Bone morphogenetic protein 7 (fragment) |

## Discussion

Non-specific immune responses are activated to limit the initial phase of parasitic invasion or infection by pathogenic micro-organisms. Parasitic invasion initiates type Th2 immune responses, characterised by the activation of Th2 lymphocytes, eosinophilia, basophilia, mast cells, and alternatively, activated macrophages (AAM). This process is accompanied by the secretion of IgE antibodies and numerous cytokines, such as IL-3, IL-4, IL-5, IL-9 IL-10, IL-13 and TGF-β. IL-13 plays a key role in regulating the anti-parasitic response [24], and is a primary factor inducing fibrosis processes in many chronic contagious and autoimmune diseases [25]. IL-13 increases the concentration of TGF-β, which leads to collagen deposition in lung and kidney tissues [26], by stimulating macrophages to produce TGF-β via the IL-13Rα2 receptor. Inhibition of IL-13Rα2 expression reduces TGF-β secretion and decreases collagen deposition in the tissues. Therefore, IL-13Rα2 is considered a feasible target molecule for therapies aimed at preventing fibrosis processes involving TGF-β [27,28]. Fibrosis is considered the final stage in the development of CKD regardless of the primary cause, and the effector cells of this process include myofibroblasts developed from tubule epithelial cells transformed during the EMT process [29,30]. During this transition, cells lose polarity, loosening their communication abilities and degrading the basement membrane. Adhesive molecules that bond both epithelial cells and the basement membrane, such as E-cadherin and integrins, are replaced by mesenchymal cell markers, such as N-cadherin, unstriated muscle α-actin, vimentin, fibronectin and collagen I. In an inflammatory environment, the EMT maintains tissue homoeostasis by inducing structural regeneration and reconstruction after harmful stress. Extinction of the inflammatory reaction results in termination of the EMT and a return to the original state. Long-term support of the EMT process leads to fibrous degeneration as well as structural and functional tissue and organ disorders [31,32]. Pleiotropic TGF-β molecules and bone morphogenetic proteins (BMPs), belonging to the transforming growth factor-β (TGF-βSF) superfamily, participate in one of the most well-known signalling pathways in the ETM process [33–35]. TGF-β plays a significant role in kidney diseases by functioning in fibrosis, inflammatory responses, apoptosis, and cell growth and diversification. Increased TGF-β levels lead to loss of the epithelial phenotype, acquisition of the mesenchymal phenotype and collagen accumulation. On the other hand, BMP-7 inhibits fibrosis, exerts anti-inflammatory effects and stimulates the regeneration of damaged kidney tissues. During ontogeny, BMP-7 has a decisive impact on the number of nephrons and the size of the organ [36]. Serine-threonine kinase receptors and cytoplasmic proteins (Smads) participate in transferring TGF-β/BMP pathway signals. By bonding with its TβRII receptor on the cell surface, TGF-β activates the TβRI receptor, which passes the signal via the phosphorylation of Smad2 and Smad3. Similarly, BMP-7 bonds its surface receptors BAMPRI and BAMPRII and phosphorylates the Smad proteins 1, 5, and 8. The phosphorylated proteins (Smads) complex with the Smad4 protein, which permeates the kernel and induces the transcription of effector genes. Smad3, induced by TGF-β stimulation, can combine with the Col1A2 gene promoter to activate the expression of type 1α2 collagen, which may accumulate in interstitial tissue and contribute to extracellular matrix (ECM) accumulation, leading to fibrous degeneration of the organ. Alternatively, TGF-β and BMP-7 expression can be controlled by extracellular signal-regulated kinases (ERKs) or mitogen-activated protein kinases (MAPKs) [37]. In experimental systems, BMP-7 recombinant protein expression or BMP-7 overexpression inhibits fibrosis during diabetic nephropathy or AKI, TGF-β-initiated EMT and E-cadherin suppression. BMP-7 manifests as a protective agent against kidney diseases by exerting an anti-inflammatory effect, reflected by the inhibition of neutrophil, monocyte and macrophage infiltration and activity as well as by repression of the expression of the proinflammatory cytokines IL-6 and IL-1β and the proinflammatory chemokines MCP1 and IL-8 [38]. BMP-7 showed anti-apoptotic activity in experiments on tubular epithelial cell (TEC) lines. The presence of α-1 antitrypsin abolishes the TGF-β/Smad3 signalling pathway and inhibits fibrosis, which indicates its therapeutic potential [39]. The BMP-7 mRNA expression in kidney bioptats of dogs with innate portal-collateral fusion was higher than that in healthy dogs. Attempts are being made to establish the causality between increased BMP-7 expression and kidney disturbances accompanying this disease, manifesting as kidney enlargement, increased glomerular filtration, polyuria and polydipsia. After pathological vessels are surgically corrected, it is important that the kidneys return to normal functioning, and the properties of BMP-7 associated with fibrosis inhibition, apoptosis and anti-inflammatory effects may be useful for accomplishing this goal, aiding in the conservative treatment of this disease [40]. Among the current concepts involving the therapeutic use of BMP-7 in kidney diseases, attention is drawn to the widespread BMP-7 receptors in various organs and the risks of side effects. Therefore, antagonists that selectively stimulate receptors associated with renal tissues are sought, and those that are present in bone tissues, for example, must be omitted[37].

## Conclusion

In summary, to the best of our knowledge, this is the first study to comprehensively analyse the urinary proteome of dogs with babesiosis, demonstrating the association of the identified proteins with the disease and indirectly confirming the occurrence of Th2 immune responses to *Babesia canis* infection. Urine Interleukin-13, bone morphogenetic protein 7, α2(1) collagen and FER tyrosine kinase are potential biomarkers of kidney damage during babesiosis in dogs that might be assigned to early renal injury; however, verifying their significance in the diagnosis and prognosis of the disease requires further study.

## Declarations of interest

The authors have declared that no competing interests exist.

## Acknowledgements

We thank Dorota Pietras-Ozga, PhD for technical assistance with the electrophoresis procedurę.

## Funding

This work was partly supported by the Polish National Science Centre (NCN) [grant numbers UMO-2016/23/N/NZ5/02576; UMO-2017/25/N/NZ5/01875]

## Role of the funding source

None to declare

